# When honest people cheat, and cheaters are honest: Cognitive control processes override our moral default

**DOI:** 10.1101/2020.01.23.907634

**Authors:** Sebastian P.H. Speer, Ale Smidts, Maarten A.S. Boksem

## Abstract

Every day, we are faced with the conflict between the temptation to cheat for financial gains and maintaining a positive image of ourselves as being a ‘good person’. While it has been proposed that cognitive control is needed to mediate this conflict between reward and our moral self-image, the exact role of cognitive control in (dis)honesty remains elusive. Here, we identify this role, by investigating the neural mechanism underlying cheating. We developed a novel task which allows for inconspicuously measuring spontaneous cheating on a trial-by-trial basis in the MRI scanner. We found that activity in the Nucleus Accumbens promotes cheating, particularly for individuals who cheat a lot, while a network consisting of Posterior Cingulate Cortex, Temporoparietal Junction and Medial Prefrontal Cortex promotes honesty, particularly in individuals who are generally honest. Finally, activity in areas associated with Cognitive Control (Anterior Cingulate Cortex and Inferior Frontal Gyrus) helped dishonest participants to be honest, whereas it promoted cheating for honest participants. Thus, our results suggest that cognitive control is not needed to be honest or dishonest per se, but that it depends on an individual’s moral default.

## Introduction

Imagine a friend sends you a link to a website where you can illegally stream recently released movies for free. Would you decide to stream the movie which you otherwise would have paid for? If so, how many movies would you stream? On a daily basis we are faced with the conflict between the temptation to violate moral standards to serve our self-interest and to uphold these moral standards, but how the brain resolves this conflict remains elusive.

When exposed to the opportunity to cheat, clearly the (financial) rewards play a crucial role in the decision-making process: the higher the reward, the more attractive the decision to cheat (Becker 1968; Allingham and Sandmo 1972). As the renowned British novelist Jonathan Gash so eloquently stated: ‘Fraud is the daughter of greed’ (Gash, 1992). In line with this sentiment, research has demonstrated that greedy people are more inclined to withhold information from others in negotiation settings (Steinel and DeDreu, 2004); they also find a variety of moral transgressions more acceptable and have engaged in such transgressions more often as compared to less greedy people (Seuntjens et al., 2019). More greedy people are also more likely to take bribes and prefer higher bribes (Seuntjens et al., 2019). Further, it has been found that neural responses in anticipation of reward, reflected in activity in the nucleus accumbens (NAcc), predict cheating behavior in a subsequent task (Abe & Greene, 2014). Collectively, these findings emphasize that higher rewards and stronger sensitivity to reward increase the likelihood of dishonesty.

Accumulating evidence from psychology, economics and neuroscience has demonstrated, however, that people care about more than only maximizing their own monetary payoff, which is reflected in the high prevalence of prosocial behaviors such as altruism and reciprocity. Through socialization, people internalize the social norms society imposes on them and use these as an internal benchmark against which they compare their behavior (Campbell 1964; Henrich et al. 2001). In the context of dishonesty, the way we view ourselves, our self-concept (Aronson 1969; Baumeister 1998; Bem 1972), may prevent us from cheating. People highly value honesty in others and also have strong beliefs in their own moral standards (Dhar & Wertenbroch, 2012). Violating one’s own moral standards will require a negative update of one’s self-concept which is highly aversive (Berthoz et al., 2006). As a consequence, people are motivated to uphold their self-concept even if it comes at the cost of potential monetary gains (Mazar, Amir, & Ariely, 2008). Hence, when given the opportunity to cheat, people are torn between the conflicting motivations to obtain a desirable monetary reward versus the long-term goal of maintaining a positive self-image.

Several lines of research have proposed that cognitive control is needed to resolve this tension between reward and self-concept (Abe & Greene, 2014; Gino, Schweitzer, Mead, & Ariely, 2011; Greene & Paxton, 2009; Maréchal, Cohn, Ugazio, & Ruff, 2017; Mead, Baumeister, Gino, Schweitzer, & Ariely, 2009). It has been found that this conflict is often settled with a compromise in which participants behave dishonestly enough to profit from the opportunity to cheat, but honestly enough to maintain a positive self-image (Mazar et al., 2008). While it is evident that cognitive control plays a crucial role in resolving this conflict, the precise nature of the role of cognitive control in moral decisions remains controversial. Two competing theories have been proposed: the ‘Will’ and the ‘Grace’ hypothesis (Greene & Paxton, 2009).

The ‘Will’ hypothesis puts forward that people are per default selfish and dishonest and that in order to be honest, deliberate cognitive control needs to be exerted. Thus, honesty is a result of the effortful resistance of temptation, similar to the cognitive control processes that allow individual to delay gratification (McClure, Laibson, Loewenstein, & Cohen, 2004). This hypothesis is supported by behavioral studies that have shown that participants who are cognitively depleted by demanding tasks are more prone to dishonest behavior (Gino et al., 2011; Mead et al., 2009). These authors argue that once our cognitive resources are taxed by prior efforts it becomes harder to resist the temptation to cheat. Similarly, sleep deprived individuals were more likely to engage in dishonest behavior not only in laboratory experiments but also in the workplace (Barnes, Schaubroeck, Huth, & Ghumman, 2011). Moreover, restraining participants from deliberate thinking through cognitive load (Welsh & Ordonez, 2014) or time pressure (Shalvi, Eldar, & Bereby-Meyer, 2012) increases cheating behavior. Collectively, these studies suggest that people automatically serve their self-interest and require cognitive control to resist the temptation to cheat in order to maintain a positive self-image.

In contrast, the ‘Grace’ hypothesis proposes that people are basically honest and require cognitive control to override their dominant honest impulses to occasionally profit from an opportunity to cheat. This is in line with the social intuitionist theory that argues that honesty is driven by moral intuitions, shaped by culture and social norms, and does not require deliberate cognitive control processes (Haidt, 2001). The hypothesis that cheating rather than honesty is a complex cognitive function demanding cognitive effort, is supported by research showing that people react faster when asked to tell the truth as compared to lying (for meta-analyses, see Suchotzki, Verschuere, Van Bockstaele, Ben-Shakhar, & Crombez, 2017; Verschuere, Köbis, Bereby-Meyer, Rand, & Shalvi, 2018). Cheating requiring cognitive capacity is also supported by findings that people cheat less when taxed by a cognitively demanding memory task as compared to a less taxing task (van’t Veer, Stel, & Van Beest, 2014). Moreover, people are less likely to deceive their opponents in an economic game when under time pressure (Carparo, 2017). In sum, these findings suggest that honesty is intuitive and cogntive control is required to override this default intuition in order to benefit from an opportunity to cheat.

In light of these evidently contradictory findings, this study aims at investigating how cognitive control resolves the conflict between external financial reward and one’s self concept and how this decision process unfolds in the brain. A better understanding about the function of cognitive control in the decision to cheat may thus help reconcile the controversy between the Will and Grace hypothesis.

In order to study how reward, self-concept and cognitive control influce cheating on a trial by trial basis, we developed an innovative task, based on the paradigm proposed by Gai and Puntoni (2017), in which participants could cheat repeatedly, deliberately and voluntarily inside the MRI scanner without suspicion of the real purpose of the task. Specifically, the advantage of this task, termed the Spot-The-Difference task, is that it allows to directly track on which trials the participants cheated, enabling us to study within subject variation in moral decisions and its neural underpinnings. Previous neuroimaging studies on cheating behavior have not been able to answer these questions as they used tasks such as the coin-flip task (Greene & Paxton, 2009; Abe & Greene, 2014), where cheating is inferred from the aggregate behavior at the end of the task thus eliminating the possibility to study trial-by-trial variation in behavior. Importantly, participants believed that the experimenter did not know that they were cheating, which is critical as participants are found to cheat less if participants believe experimenters can observe the true outcome (Gneezy, Kajackaite, & Sobel, 2018). The Spot-the-Difference paradigm is therefore the first behavioural paradigm to assess cheating behaviour inconspicuously on a trial-by-trial basis enabling us to study individual differences in neurocognitive processes underlying cheating behaviour while also being sensitive to within subject variation.

In our analysis, we first conducted an exploratory whole brain analysis to identify the brain networks underlying the decision to cheat or to be honest. We first identified the brain networks engaged when exposed to the opportunity to cheat and when making the decision to cheat or to be honest. To reduce the reverse inference problem (Poldrack, 2006), we then assessed the neural overlap between our results and meta-analytically derived maps associated with, respectively, reward, self-concept and cognitive control from Neurosynth (Yarkoni et al., 2011). Subsequently, we used the ROIs obtained from this conjunction analysis to conduct a trial-by-trial analysis to study the neural mechanisms underlying within-subject variation in cheating behavior and also to explore functional connectivity between the resulting networks of regions. To test the generalizability and replicability of our results, we then used cross-validation to explore whether we can use neural activation to predict unseen trials, and functional connectivity patterns to distinguish between cheaters and honest participants.

## Method

### Participants

The reported analyses are based on 40 participants (30 females; age 18-35 years; *M* = 23.7 *SD* = 3.2) recruited from an online community for university students, where students can sign up for experiments. An intial screening interview ensured that all participants were right-handed with normal or corrected to normal vision, spoke English fluently, were not on any psychoactive medication influencing cognitive function and had no record of neurological or psychiatric illness. The study was approved by the university Ethics Committees and was conducted according to the Declaration of Helsinki.

### Task and Stimuli

#### Spot-The-Difference Task

In the Spot-The-Difference task, participants were presented with pairs of images and were told that there were always three differences between the image pairs. Differences consisted of objects that were added to or removed from an image, or objects that differed in colour between images. However, images could actually contain one, two, or three differences. Participants were asked to find three differences between the images. Because reward (see below) was contingent on participants *reporting* that they had found all three differences, without having to point them out, this design encouraged cheating behavior (i.e., reporting having found all three, even when objectively fewer than three differences were present in the images).

Participants were told that the purpose of the study was to investigate the underlying neural mechanisms of visual search for marketing purposes such as searching for a product in an assortment or information on a webpage. In order to increase credibility of this cover story a simple visual search task was added at the beginning of the experiment (see Appendix 1), which was also performed in the scanner while participants were undergoing localizer scans. Further, participants were instructed that the neurocognitive effect of motivation, elicited by monetary reward, on speed and accuracy of visual search was investigated. Although participants were told that there were three differences in all trials, in 25% of the trials there were only two differences and in 25% there was only one difference. All stimuli were standardized in size and were presented on a white background on a computer screen. The ratio of 50% - 50% (three differences vs less than three differences) was chosen based on the results of pilot studies that indicated this ratio to be optimal in reducing suspicion that the pairs did not always contain three differences.

Trials were further categorized into normal (50%), hard (25%) and very hard trials (25%), for which participants could receive 5cts, 20cts, and 40cts, respectively. All of the trials with three differences (the filler trials) were categorized as normal trials, whereas trials with less than three differences (the trials of interest) were randomly categorized as hard or very hard trials. Consequently, the reward was independent of the number of differences in the image pair for the trials of interest, which is important in order to be able to disentangle the effects of reward and cheating magnitude (the actual number of differences) on cheating behavior. The different levels of difficulty were added to reduce suspicion about the real purpose of task. It was assumed that if trials are labeled as hard or very hard it would be more credible to the participant that the image pair actually contained three differences, but they were just too hard to spot. In addition, levels of difficulty were introduced to eliminate possible demand effects: we wanted participants to cheat for monetary reward and not to prevent seeming incompetent, which may be associated with different underlying neural mechanisms and consequently confound the analysis.

To further reduce suspicion about the purpose of the study, approximately 10% of all trials were point-and-click trials. In these trials, participants had to click on the location in the images where they spotted the differences using a joy-stick. As a consequence, cheating was not possible on the point-and-click trials. Participants always knew prior to the start of a trial whether it was a point-and-click trial indicated by a screen requesting participants to click on the image. This ensured that participants would not refrain from cheating on all other trials, while still reducing the suspicion about the real purpose of the study. Participants were told that only 10% of trials were point-and-click trials because it would take too much time to point out the differences for every pair. Further, participants were instructed that excessive movement by manipulating the joystick would interfere with the brain signal. In sum, there were 144 regular trials (of which 72 cheatable trials) and 12 point-and-click trials. The maximum amount of money earned, in case a participant cheated on all cheatable trials was approximately 35 Euros, whereas in case a participant would not cheat at all he or she would earn approximately 7.50 Euros. To be fair to all participants, after completion of the full study, participants were debriefed and they were all paid out the maximum amount, irrespective of their actual cheating behavior.

Each trial started with a fixation cross which was presented for a variable amount of time between 1-3s (see Figure 1). Subsequently, the Level of Difficulty screen was presented for 2 seconds informing the participants about the level of difficulty of the upcoming trial. This screen also displayed how much money could be earned on that trial. As a result, participants were constantly aware of the potential gains of cheating. Next, an image pair was presented for 6s, a length determined by the behavioral pilots, and participants engaged in the visual search. Afterwards, the participants were asked whether they spotted all three differences (yes/no response). On this decision phase screen, again the potential reward for this trial was presented, in order to make the reward more salient and increase cheating behavior. After 3s, the response phase started in which participants’ responses were recorded. In the decision phase and the response phase the current balance was also shown, which was done to demonstrate to the participants that if they stated that they had found the three differences, their current balance increased immediately. It was assumed that this direct noticeable effect of behavior on the increase of the current balance, would further motivate participants to cheat.

**Figure 1.**
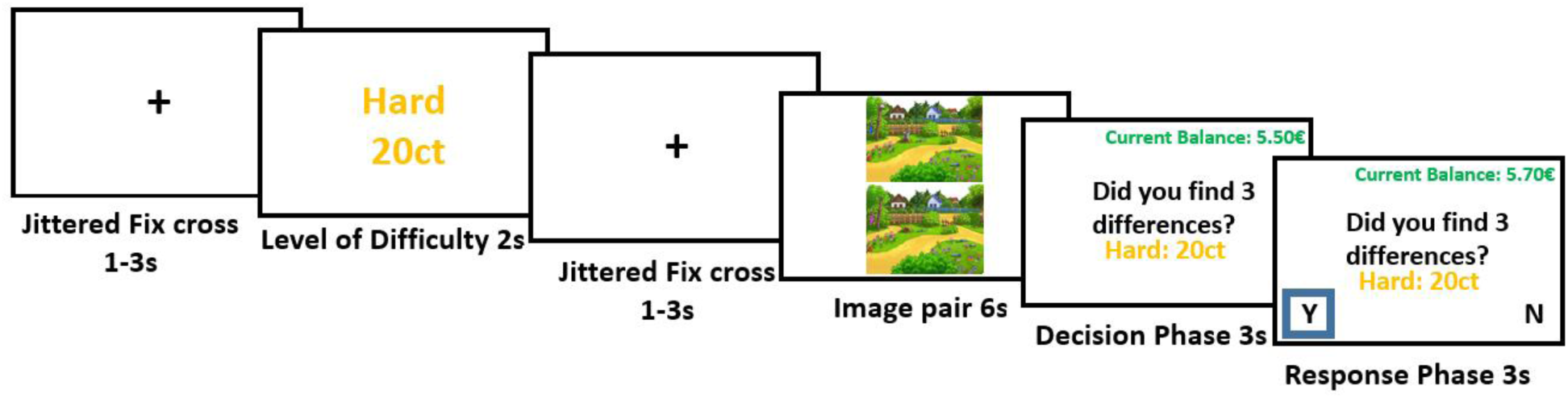
One trial of the Spot-The-Differences paradigm. Participants view a screen indicating the difficulty and value of the trial, then the image pair appears for six seconds and then participants have to indicate whether or not they spotted all three differences.

The decision phase and response phase were separated to isolate the decision from motor responses. This was important for the fMRI analysis as we wanted to isolate the neural mechanisms underlying decision making from possible neural confounds related to button presses. Besides that, the buttons corresponding to “Yes” and “No” were switched across trials to further reduce confounding effects and to reduce the response bias for the dominant hand. Once the participants responded, the choice was highlighted by a blue box for 500ms to indicate that the response was recorded and the trial ended. If no response was made, the trial ended after 3s. In addition, there were five practice trials, in which participants could get acquainted with the task. Stimulus presentation and data acquisition was performed using Presentation® software (Version 18.0, Neurobehavioral Systems, Inc., Berkeley, CA, www.neurobs.com).

The main advantage of our experimental design is that it allowed tracking on which trials the participants cheated. As we knew how many differences there are in each image pair, we knew precisely whether the participants cheated or not. Further, by varying the number of differences, this design enabled us to assess the magnitude of cheating (i.e, cheating when only 1 vs 2 differences were found). It is therefore the first behavioural paradigm that allowed to assess cheating behaviour inconspicuously on a trial-by-trial basis in the scanner.

#### Stimuli

Stimuli for the task consisted of 144 Spot-The-Difference image pairs that were downloaded from the Internet. Cartoon images of landscapes containing several objects were selected, to make them engaging and challenging enough for the participants. Landscapes were chosen as they generally satisfied the necessary criterion of containing several different objects. The stimuli consist of pairs of images that are identical apart from a certain number (1-3) of differences that were created using Adobe Photoshop. Differences consisted of objects added to or removed from the landscape picture or changed colors of objects. Differences were fully randomized across all pairs of images, which means that all image pairs could be presented with either one, two or three differences. To make sure that participants would be able to find the differences between the images in a reasonable amount of time, we ran a pilot study on Amazon’s Mechanical Turk (N=205) to test the difficulty to spot the differences between the images and to determine the optimal duration of picture presentation (see Appendix 2).

#### Experimental procedure

Before the experiment started, participants were introduced to the cover story, the tasks and the scanner environment and they signed the informed consent form. They were then informed about and checked on the safety requirements for MRI scanning and completed practice trials for both visual search tasks outside of the scanner. Subsequently they were guided into the scanner and completed the simple visual search task (5 min) followed by the Spot-The-Difference task which took approximately 45 minutes. After completing the two tasks in the scanner, participants were taken to a separate room in absence of the experimenter and filled-in a short questionnaire including questions about their thoughts on the purpose of the task.

#### FMRI acquisition

The functional magnetic resonance images were collected using a 3T Siemens Verio MRI system. Functional scans were acquired by a T2*-weighted gradient-echo, echo-planar pulse sequence in descending interleaved order (3.0 mm slice thickness, 3.0 × 3.0 mm in-plane resolution, 64 × 64 voxels per slice, flip angle = 75°). TE was 30ms and TR was 2030ms. A T1-weighted image was acquired for anatomical reference (1.0 × 0.5 × 0.5 mm resolution, 192 sagittal slices, flip angle = 9°, TE = 2.26ms, TR = 1900ms).

### fMRI analysis

#### Preprocessing

The fMRI data was preprocessed using fMRIPrep version 1.0.8, a Nipype based tool (Gorgolewski et al., 2011). The reason for choosing fMRIPrep was that it addresses the challenge of robust and reproducible preprocessing as it automatically adapts a best-in-breed workflow to virtually any dataset, enabling high quality preprocessing without the need of manual intervention (Esteban et al., 2019). Each T1w volume was corrected for intensity non-uniformity (INU) and skull-stripped. Spatial normalization to the ICBM 152 Nonlinear Asymmetrical template version 2009c (Esteban et al., 2016) was performed through nonlinear registration, using brain-extracted versions of both T1w volume and template. Brain tissue segmentation of cerebrospinal fluid (CSF), white-matter (WM) and gray-matter (GM) was performed on the brain-extracted T1w. Fieldmap distortion correction was performed by co-registering the functional image to the same-subject T1w image with intensity inverted (Caballero-Gaudes & Reynolds, 2017) constrained with an average fieldmap template (Tustison et al., 2010). This was followed by co-registration to the corresponding T1w using boundary based registration (Smith, 2002) with 9 degrees of freedom. Motion correcting transformations, field distortion correcting warp, BOLD-to-T1w transformation and T1wto-template (MNI) warp were concatenated and applied in a single step using Lanczos interpolation. Physiological noise regressors were extracted applying CompCor (Cox & Hyde, 1997).

Principal components were estimated for the two CompCor variants: temporal (tCompCor) and anatomical (aCompCor). Six tCompCor components were then calculated including only the top 5% variable voxels within that subcortical mask. For aCompCor, six components were calculated within the intersection of the subcortical mask and the union of CSF and WM masks calculated in T1w space, after their projection to the native space of each functional run. Frame-wise displacement (Treiber et al., 2016) was calculated for each functional run using the implementation of Nipype. For more details of the pipeline see https://fmriprep.readthedocs.io/en/latest/workflows.html.

#### Statistical analyses

For each participant we estimated a general linear model (GLM) using regressors for onsets of the decision phase for cheated trials, honest trials, cheatable trials (trials with less than three differences) and non-cheatable trials (trials with three differences). The duration of the epoch for the decision phase was three seconds and the beginning of the decision phase was used as onset times. The decision phase was used as it provides all the necessary information to make the decision and is free of brain activity related to motor responses. In addition, regressors were added for the onsets of the Level of difficulty phase with a separate regressor for each level of reward. For the level of difficulty trial the duration was two seconds. This phase was used to test whether participants are indeed sensitive to differences in potential gain as it provided information about the possible reward without any moral conflict. Besides that, in order to ensure that there were no significant differences in engagement or motivation in the Spot-The-Difference task between conditions or subjects, regressors were added for the onsets of the visual search phase in which the image pairs were presented on the screen. The duration of the visual search phase was six seconds (see *Figure 1*). Lastly, regressors for the button presses were added. Average background, white matter and cerebrospinal fluid (CSF) signal, framewise displacement, six head motion regressors and six aCompCor regressors, all obtained from fMRIprep, were entered as regressors of no interest. All regressors were convolved with the canonical hemodynamic response function. A smoothing kernel of 5 mm (FWHW) was applied. Linear contrasts were computed between honest and cheating decisions and between cheatable and non-cheatable trials. These contrasts were then subjected to a random effects analysis to compute main effects (one sample t-test), and to regression analyses with behavioral data (i.e., total amount of cheating for each participant) as regressors.

#### Cheatable vs. Non-cheatable trials

To identify the neural correlates associated with the opportunity to cheat, we contrasted the neural activation during cheatable trials (trials with less than three differences), against activation in non-cheatable trials (trials with three differences) in both directions. Subsequently, using the contrast images obtained for each subject, one sample t-tests were conducted on the group level to explore the average effect of being exposed to the opportunity to cheat across participants. We also added the cheat count, which is a measure how often each participant cheated in total on the Spot-The-Difference task, as a group level covariate to explore whether there are individual differences in the neural mechanisms when exposed to the opportunity to cheat, between individuals who cheat a lot vs. those who rarely cheat. The threshold applied to the group level statistical maps was a voxel-wise false discovery rate of p < 0.05 (FDR) to correct for multiple comparisons. Clusters of activation resulting from the thresholding were characterized in terms of their peak voxels in the MNI coordinate space.

#### Honest decisions vs. Cheating

To explore the neural mechanisms underlying the decision to cheat, we contrasted neural activation in the decision phase on trials on which participants cheated against honest trials in both directions. For each of these contrasts we then conducted one sample t-tests on the group level to explore the average effects of each of these contrasts across participants. In addition, we also entered the total cheat count for each participant as covariate on the group level to investigate the correlation between behavior and neural activation in the contrasts of interest. Based on the resulting beta images, second-level random-effects group contrast maps were then created in both directions (i.e., positive and negative correlation between activation and cheat count). The threshold applied to the group level statistical maps was a voxel-wise false discovery rate of p < 0.05 (FDR) to correct for multiple comparisons. Clusters of activation resulting from the thresholding were characterized in terms of their peak voxels in the MNI coordinate space. Due to the fact that participants engaged in spontaneous, voluntary and deliberate cheating, the proportion of cheated and honest trials was not balanced for most of the participants. To account for possible confounding statistical effects of this imbalance, we under sampled the majority class for each participant to create a perfect balance when estimating the contrasts (Liu, Wu, & Zhou, 2008).

#### Single trial activation estimation

An important contribution of our task is that it allows us to assess cheating behavior on a trial-by trial-basis. That is, we are able to assess why a person who is generally honest, decides to cheat on some trials, and why a cheater might refrain from cheating on some occasions. To explore which neural mechanisms underlie this within subject variability we extracted the neural activation from the ROIs identified in the analyses described above during decision making for each trial for each subject. These trial-by-trial activations could then be fed into multilevel models to explore which neural mechanisms may explain within subject variability.

To obtain single trial neural activations for the trial-by-trial multilevel models, individual time series were modeled using a double γ hemodynamic response function in a single trial GLM design using FSL’s FEAT. Specifically, one GLM fitted a hemodynamic response function (HRF) for each trial, following the Least-Squares all (LSA) approach (Mumford, Turner, Ashby, & Poldrack, 2012), using the decision phase and level of difficulty phase of each trial, resulting in parameter estimates of sustained activations for each trial for each participant. The resulting β-values were converted to t-values (Misaki, Kim, Bandettini, & Kriegeskorte, 2010), resulting in a whole-brain map of t-values for each trial. The duration of the epoch for the decision phase was 3 seconds and 2 seconds for the level of difficulty phase. As for the previous analyses, average background, white matter and CSF signal, framewise displacement, six head motion regressors and six aCompCor regressors, all obtained from fMRIprep, were entered as regressors of no interest. All regressors were convolved with the canonical hemodynamic response function. Mulilevel modelling was conducted with custom R scripts in combination with the ‘lme4’ package for linear mixed-effects models (Bates et al., 2015) and the ‘glmmlasso’ package for variable selection for generalized linear mixed models by L1-penalized estimation (Groll & Tutz, 2014). fMRI analyses were conducted using custom Python scripts, which will be made publicly available.

#### Beta-series correlations

In order to further explore how the different areas resulting from the different contrasts described above interact with each other during decisions to cheat, we investigated the functional connectivity between these areas during the decision phase of the Spot-The-Difference Task. To avoid the problem of activation-induced correlations we implemented beta-series correlations (Rissman, Gazzaley, & D’Esposito, 2004). We used the single trial activations obtained as explained above by fitting a model that includes a separate regressor for each trial. We then correlated the parameter estimates from these regressors (the “beta series”) for honest decisions and cheated decisions separately between all the regions found to be significantly related to our contrast of interest, in order to examine the degree to which they show similar trial-by-trial activations, as is expected when these regions were functionally connected. The beta-series model is particularly useful in event-related fMRI studies where the spacing between trials is relatively long (more than 8-10 seconds), which is the case in our paradigm (Poldrack, Mumford, & Nichols, 2011). After obtaining the correlation matrix for each of the participants for honest and cheated decisions, we then also correlated the functional connectivity between each of the regions with the cheat count (individual differences in total cheating) in order to examine how functional connectivity differed for cheaters and more honest participants. To compare functional connectivity between honest and cheated decisions, correlations were transformed to z-values using the Fisher r-to-z transformation. Significance was estimated by means of permutation testing where the cheat count was randomly shuffled at each iteration (N=5000). The resulting empirical p-values were then corrected for multiple comparisons at FDR < 0.05.

## Results

### Behavioral results

Large individual differences in the total amount of cheating were observed (Mean= 25%, Median=14%, SD=26%; see Figure 2): some participants cheated only on one or two trials (20% of participants), whereas others only missed one or two opportunities to cheat (5 %). To assess suspicion about the real purpose of the study participants were asked what the goal of the experiment was. Participants mentioned marketing research, consumer decision-making, neuromarketing and visual search as our general cover story suggested that visual search is important for quickly locating one’s favorite brand or product in a supermarket. Importantly, none of the participants mentioned dishonesty, moral decision making or related concepts, which indicates that none of the participants were suspicious of the real goal of the study.

**Figure 2.**
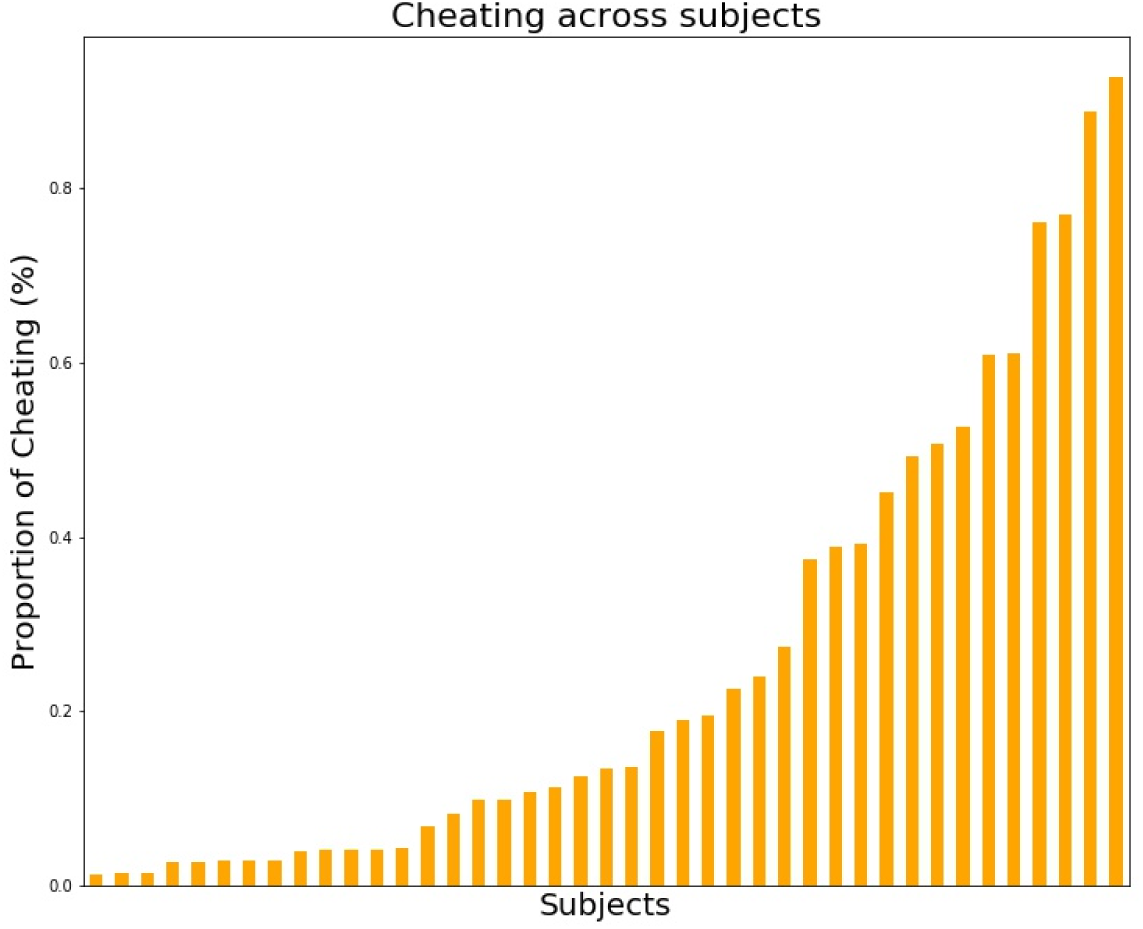
Individual differences in proportion of cheating (%) on the Spot-The-Difference task. N = 40.

We also explored how the task characteristics of the Spot-The-Difference task influenced cheating behavior. Given the nested structure of our data (trials within different number of differences and rewards within participants), we conducted a multilevel analysis for our behavioral data. This analysis was conducted for the cheatable trials only, so all trials with three differences between the images were removed. The dependent variable was the binary response (cheating vs. honest) with a logit link (cheating = 1, honest = 0). The number of differences and level of reward served as trial level predictors. The model allowed for random intercepts and random slopes within participants. This analysis revealed a significant effect of the number of differences (excluding three differences trials) on cheating behavior (*b*=2.20, *SE*=0.46, *z*=4.744, *p*<0.001). This shows that participants cheated more when the crime is smaller (that is, they indicated to have found three differences more often when they had actually found two differences as compared to when they found only one). Specifically, when there were two differences, participants cheated on 35% of the trials, whereas participants only cheated on 15% of trials with only one difference (t=3.25, p=0.002). No effect of reward magnitude on cheating behavior was observed, and no significant interaction effects between number of differences and reward were found. We also tested for possible fatigue or habituation effects by using trial number as a trial level predictor to see whether cheating behavior increased or decreased over the course of the experiment. No time effects were found.

### Neural mechanisms associated with the opportunity to cheat

#### Whole-brain analysis

As a first step of our fMRI analysis we explored the neural activation in response to the opportunity to cheat. In order to do so, we contrasted neural activity during trials in which participants had the opportunity to cheat against trials in which they did not have this opportunity (see Methods for details). To explore whether there are individual differences in the neural response to this opportunity, participants’ cheat count was added as a group level covariate. The whole brain analysis revealed that more honest participants (compared to those who cheated more) exhibited greater activation in the posterior cingulate cortex (PCC), the medial prefrontal cortex (MPFC) and the bilateral temporoparietal junction (TPJ) when exposed to the opportunity to cheat (p_FDR_<0.05; see Figure 3A and Appendix 4 for table with clusters).

**Figure 3.**
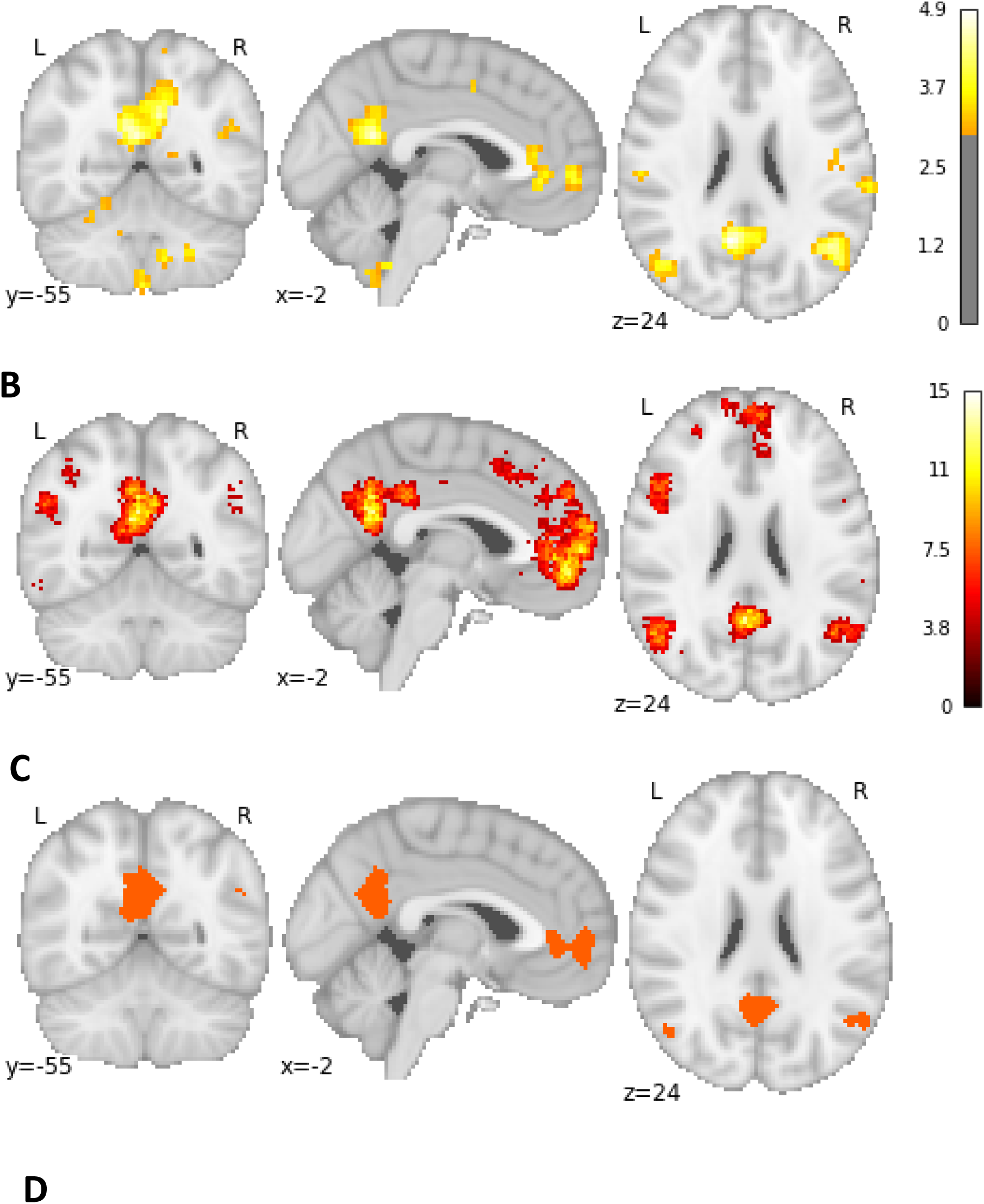
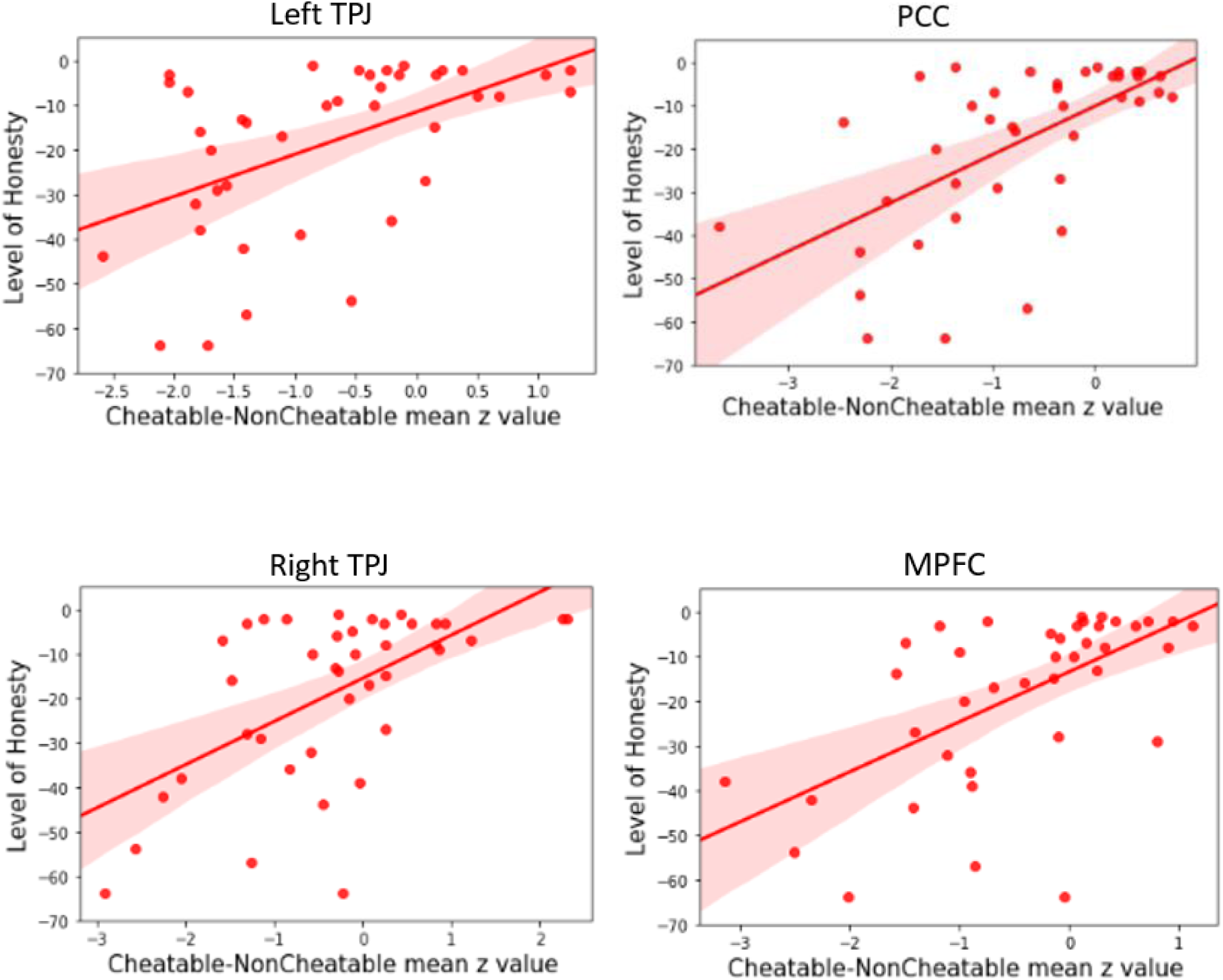
Honest participants engage their self-referential thinking network more strongly than cheaters when exposed to the opportunity to cheat. (A) More honest participants exhibit great activation in the PCC, MPFC and bilateral TPJs when exposed to the opportunity to cheat. (B) Self-referential thinking mask obtained from Neurosynth. (C) Neural overlap between group level results for cheatable vs. non-cheatable trials correlated with cheat count and the self-referential thinking mask obtained from Neurosynth (D) The correlation between the level of honesty (reversed cheat count) and neural activation when participants were exposed to the opportunity to cheat as contrasted to no opportunity trials, for the PCC, bilateral TPJs and the MPFC, respectively (using ROIs obtained from the conjunction analysis).

As the activated network in our group-level results highly resembled the self-referential thinking network, we conducted a conjunction analysis with a meta-analytically derived self-referential thinking mask obtained from Neurosynth, FDR corrected for multiple comparisons at p<0.01 (using the term “self referential”; Wager, Nichols, Van Essen, Poldrack, & Yarkoni, 2011, See Figure 3B and Appendix 3) to test whether there is indeed neural overlap. Neural overlap was found in the PCC (overlap (mm^3^) = 4600), in the MPFC (overlap (mm^3^) = 4072), in the right TPJ (overlap (mm^3^) = 869) and the left TPJ (overlap (mm^3^) = 608), see Figure 3C.

#### ROI analysis

To test whether the relationship between activity in the self-referential thinking network and level of honesty during the opportunity to cheat remains when using the ROIs obtained from the conjunction analysis, we extracted the mean activations from the contrast maps obtained above (Cheatable > NonCheatable) from each of the regions identified in the conjunction analysis (see Figure 3C). We found a positive correlation between level of honesty (reverse cheat count) and the mean activity in the PCC (*r*=0.62, *p_adj_ < 0.001;* adjusted for multiple comparisons using FDR at p<0.05), the MPFC (*r*=0.48, *p_adj_ < 0.01*), the left TPJ (*r*=0.51, *p_adj_ < 0.001*), the right TPJ (*r*=0.59, *p_adj_ < 0.001*) and the MPFC (*r*=0.59, *p_adj_ < 0.001*). This illustrates that honest participants engage their self-referential thinking network more strongly than less honest participants when exposed to the opportunity to cheat (see Figure 3D).

### Neural mechanisms underlying the decision to cheat

#### Whole-brain analysis

Next, we explored which neural mechanisms underlie the decision to cheat or not, when given the opportunity. To answer this question, we contrasted the neural activation of trials where participants had the opportunity to cheat but decided to be honest, against trials on which participants decided to cheat. As before, to explore whether there are individual differences in the neural processes underlying honest as compared to dishonest decisions, participants’ cheat count was added as a group level covariate.

We found that participants who overall cheated more, showed higher activity in the anterior cingulate cortex (ACC) and the inferior frontal gyrus (IFG) when they made the decision to be honest (p<.001, uncorrected; see Figure 4A). Stated differently, cheaters engage their ACC and IFG more than honest participants when refraining from cheating. As the activated network in our group-level results highly resembled regions within the cognitive control network, we conducted a conjunction analysis with a meta-analytically derived cognitive control mask obtained from Neurosynth (Wager, Nichols, Van Essen, Poldrack, & Yarkoni, 2011, See Figure 4B and Appendix 3) to test whether there is indeed neural overlap. Neural overlap was found in the ACC (overlap (mm^3^) = 168) and in the left IFG (overlap (mm^3^) = 1256), see Figure 4C.

**Figure 4.**
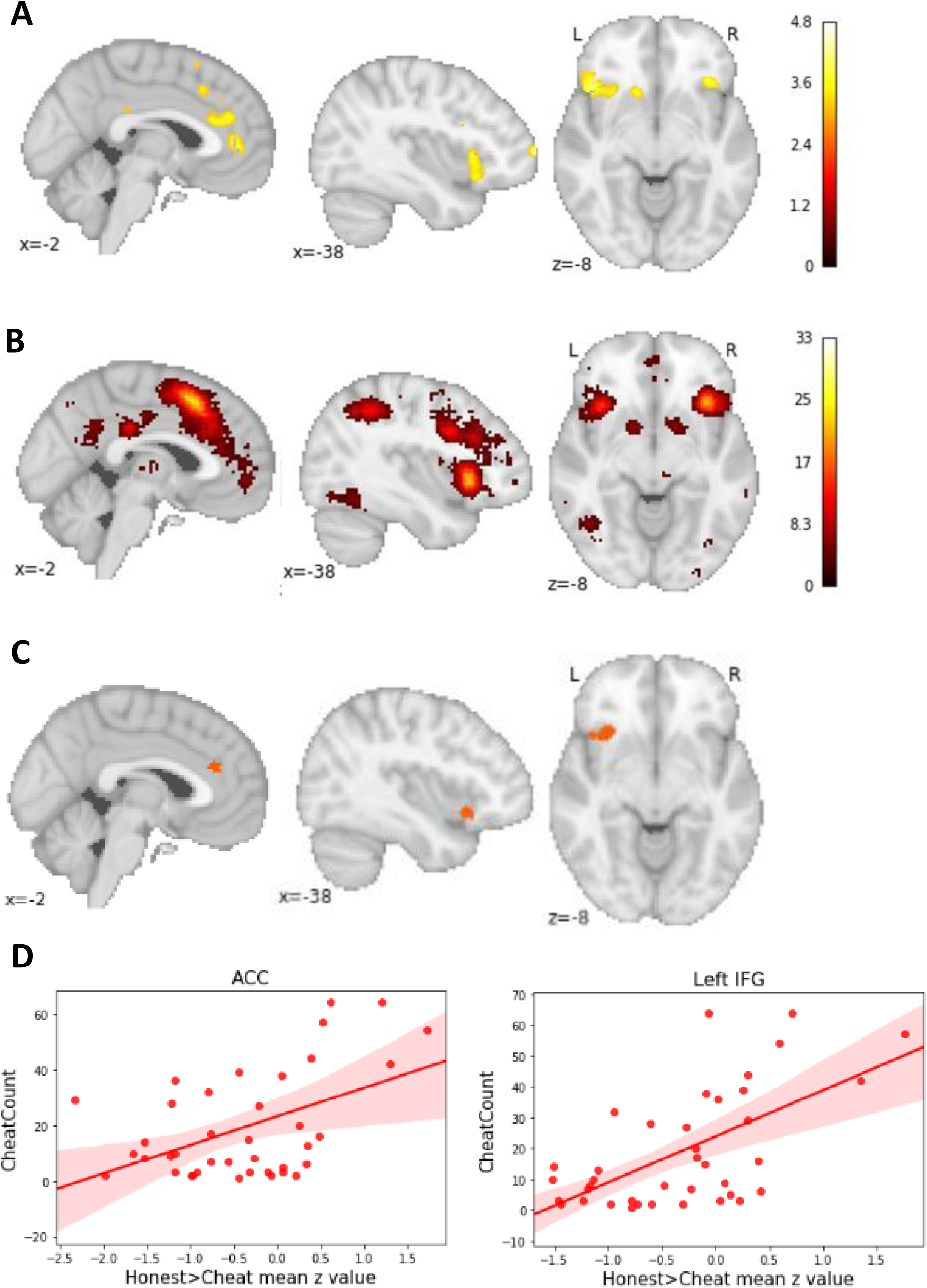
Cheaters exhibit higher activation in the ACC and left IFG when deciding to be honest. (A) Participants who cheat more, exhibit higher activation in the ACC and left IFG when making the decision to be honest. (B) Cognitive Control network derived from Neurosynth.(C) Neural overlap between group level results for honest vs. cheated trials correlated with cheat count and the cognitive control mask obtained from Neurosynth in the Left IFG and ACC. (D) The correlation between cheat count and neural activation when participants decided to be honest as contrasted to cheating decision, for the left IFG and the ACC.

#### ROI-analysis

To assess whether the association between activity in the cognitive control regions and cheatcount during honest decisions remains when using the ROIs resulting from the conjunction analysis, an ROI analysis was conducted in which we extracted the mean activation from the contrasts maps obtained in the previous analysis (honest>cheat) from each of the regions obtained from the conjunction analysis above (see Figure 4C). We found a strong positive correlation between cheat count and mean activity in the left IFG (*r*=0.61, *p_adj_ < 0.001*) and the ACC (*r*=0.49, *p_adj_ < 0.001*). These results confirm findings from the whole brain analysis and show that participants who cheat a lot engage their cognitive control network more strongly when they decide to be honest than honest participants do.

### Neural correlates of the sensitivity to reward are associated with cheating

#### Level-of-Difficulty Phase

Although we did not find any effects of reward on cheating on the behavioral level, we did want to test whether the participants responded to the reward on the neural level, as previous research has eluded to the relevance of reward anticipation in explaining individual differences in cheating (Abe & Greene, 2014, Seuntjes et al., 2019). Here, we investigated whether participants were motivated by the possible rewards that could be obtained at each trial and whether participants differentiated between the different magnitudes, 5ct, 20ct and 40ct, of reward on the neural level. We conducted a parametric modulation analysis where we used the onsets of the level of difficulty phase of each trial and added the magnitude of reward at each trial as a parametric modulator on the first level. The analysis revealed that the magnitude of reward modulated the activity in the bilateral Nacc significantly (p_FDR_<0.05; see Figure 5A and Appendix 6 for table with clusters).

**Figure 5.**
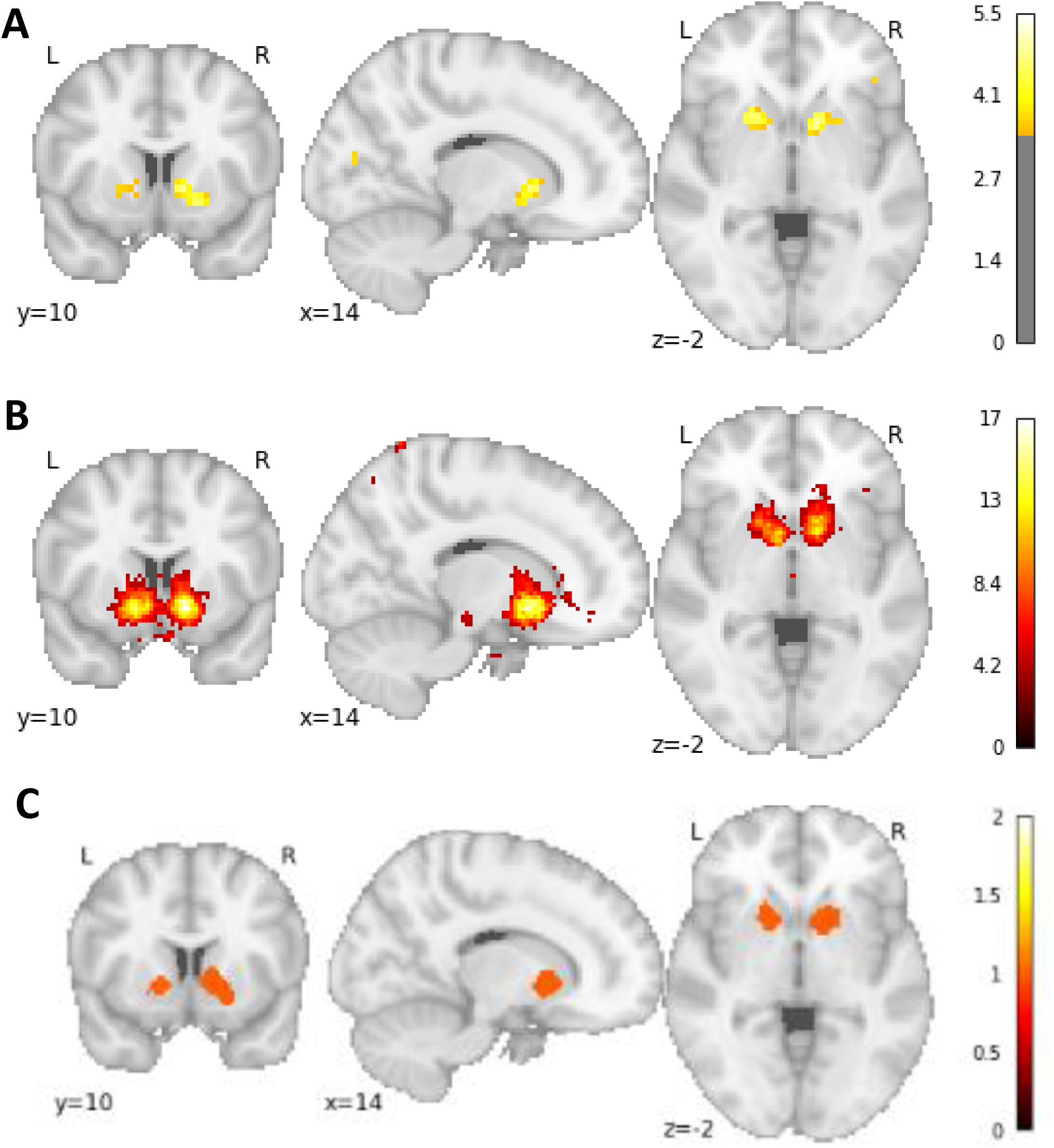
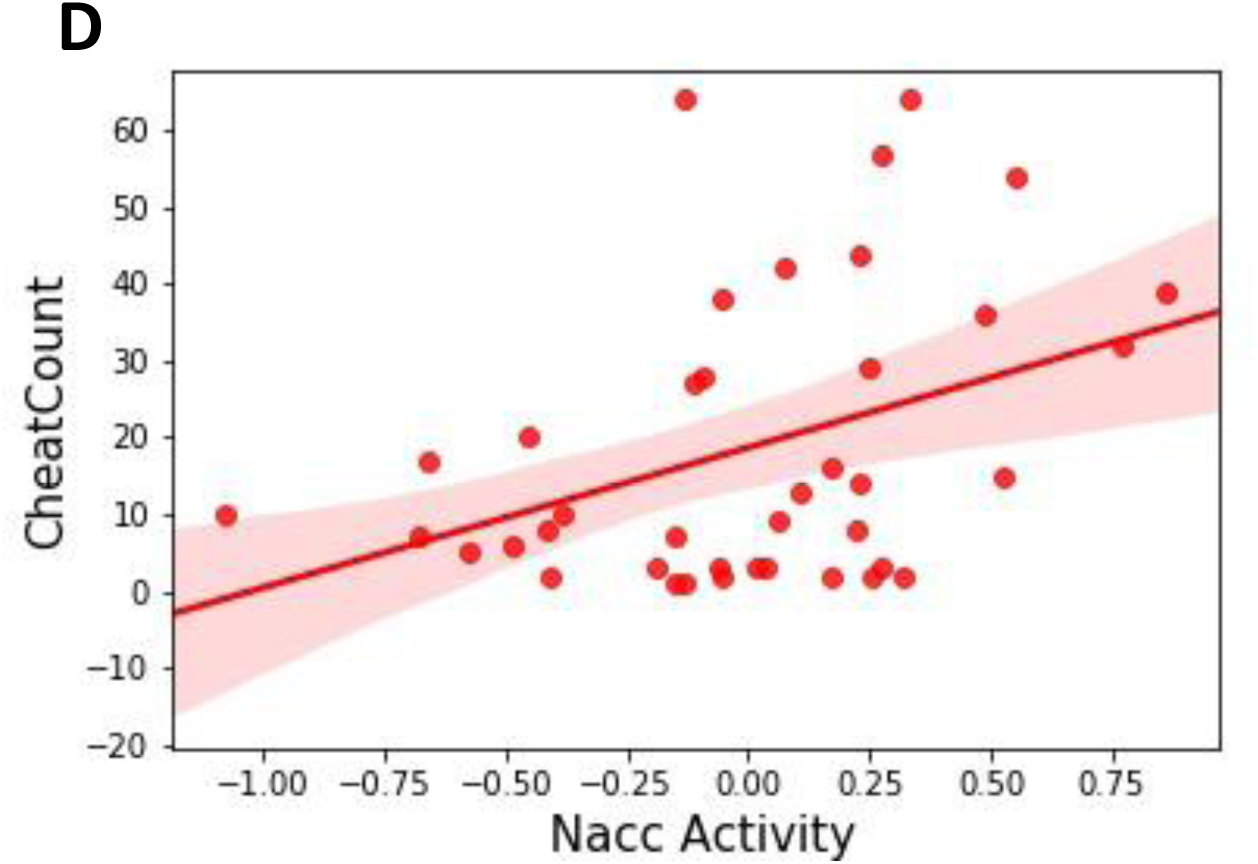
Cheaters exhibit higher activity in the Nacc when making (dis)honest decisions. (A) The left and right nucleus accumbens are parametrically modulated by the magnitude of reward. (B) Reward network derived from Neurosynth.(C) Neural Overlap between the parametric modulation analysis of the magnitude of reward and the reward anticipation network derived from Neurosynth. (D) Mean Nacc activity during the decision phase predicts cheat count.

As the Nacc is well known for its role in processing the anticipation of reward (Ballard & Knutson, 2009; Knutson, Adams, Fong, & Hommer, 2001; Oldham et al., 2018) this suggests that the participants were indeed motivated by the potential rewards presented at the beginning of the trial. Further, differences between levels of magnitude were reflected in different levels of activity in the Nacc, suggesting that participants were indeed differentiating between the different reward magnitudes.

In addition, significant activation was found in the left cuneus which is involved in basic visual processing (Vanni, Tanskanen, Seppä, Uutela, & Hari, 2001). This may reflect neural activation in response to the different visual information associated with the different levels of reward, such as different colors (green for the normal trials, orange for hard trials and red for very hard trials) and text. Alternatively, it may also reflect increased visual attention when the reward is higher.

As the activated network in our second level results highly resembled the reward anticipation network, we conducted a conjunction analysis with a meta-analytically derived reward anticipation mask obtained from Neurosynth with false discovery rate (FDR) corrected for multilple comparisons at p<0.01 (Wager, Nichols, Van Essen, Poldrack, & Yarkoni, 2011, See Figure 5B and Appendix 3) to test whether there is indeed neural overlap. Neural overlap was found in the right Nacc (overlap (mm^3^) = 2040) and left Nacc (overlap (mm^3^) = 840), see Figure 5C. We also conducted an additional second level analysis, in which we added the cheat count as a covariate, in order to explore whether reward sensitivity in the Level-of-Difficulty phase differed between subjects. However, no significant differences were observed, indicating that participants were equally sensitive to the rewards, independent of how often they cheated.

#### Decision Phase

To explore how the effect of reward anticipation, as represented by activity in the Nacc, on cheating differs for cheaters and more honest participants, we then used the ROIs derived from the conjunction analysis between our parametric modulation analysis and the Neurosynth map for reward (see Figure 5C) and regressed mean Nacc activity per subject during the anticipation and decision phase against the cheat count. This analysis revealed that average Nacc activity significantly predicted cheat count (*b*=18.29, *SE*=7.01, *p*<0.05, see Figure 5D) during the decision phase, whereas no significant effect was found during the anticipation phase (*b*=-8.89, *SE*=14.2, *p*=0.54). This suggests that participants are equally sensitive to reward during the anticipation phase when there is no moral conflict, however, when making the decision to cheat (or be honest), participants who cheat seem to be driven more strongly by anticipation of reward.

### Investigating within-subject variation in cheating: Trial-by trial analysis

In order to further explore how self-concept, reward and cognitive control influence decisions to cheat, we conducted a trial-by-trial analysis, which allowed us to explore within-subject variability. Stated differently, this analysis was aimed at investigating the neural mechanisms that determine why the same person may cheat on some occasions and remain honest on others. As a first step, we extracted average trial-by-trial activation from individual regions within the reward, cognitive control and self-referential thinking, respectively, where we used the conjunction between our second level results and the Neurosynth maps (see Figures 3C, 4C & 5C), resulting in one data matrix where the rows represent trials and the columns represent the regions of interest. Given the nested structure of our data (trials within different number of differences and rewards within participants) we then conducted a multilevel analysis for each of the networks (self-referential thinking, cognitive control and reward). The dependent variable was the binary response with a logit link (cheating = 1, honest = 0). The averaged activity within the obtained regions of interest served as trial level predictors, whereas the cheat count served as a subject level predictor. The models allowed for random intercepts and random slopes within participants.

#### Assessing the relative importance of the networks

To investigate which of the networks is most important in predicting cheating on a trial-by-trial level, we performed variable selection for generalized linear mixed models by means of L1-penalized estimation. This was implemented using the ‘glmmlasso’ package in R, which implements a gradient ascent that allows to maximize the penalized log-likelihood yielding models with reduced complexity (Groll & Tutz, 2014). The lasso regression adds a penalty term to the equation which shrinks coefficients in the model to zero and thus reduces complexity of the model and multicollinearity of predictors (Tibshirani, 1996). In this way it also selects the most important predictors in the model. This analysis revealed that the ACC (*b* = 0.13, *SE*=0.06, *p*=0.02), the left IFG (*b* = 0.42, *SE*=0.06, *p*<0.001), the cheat count (*b* = 1.59, *SE*=0.07, *p*<0.001) and the interaction effect between the left IFG and the cheat count (*b* = −0.38, *SE*=0.06, *p*<0.001), were most important in predicting cheating. These results suggest that the cognitive control network is most important in predicting cheating on the trial level. Inspecting the plot of the interaction effect (see Figure 6), we see that for participants who cheat a lot (light blue lines), higher levels in the left IFG are associated with lower probabilities of cheating, whereas for more honest participants (dark blue lines), higher activity in the left IFG is associated with higher probability of cheating. These findings suggest that the effect of the left IFG on cheating depends on whether a participant has the general tendency to cheat or to be honest.

**Figure 6.**
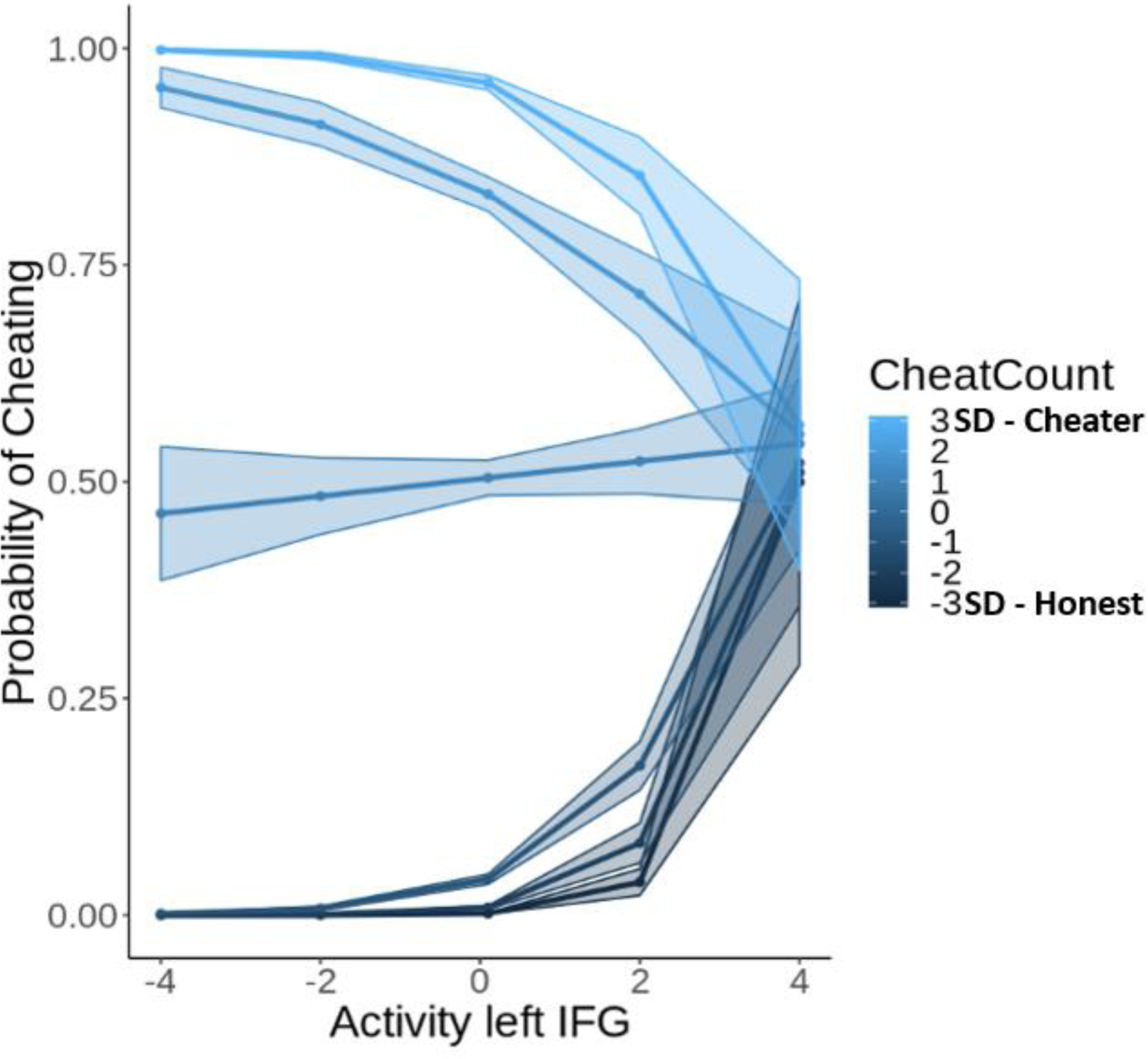
Interaction effect between cheat count and the left IFG in predicting the probability of cheating. The lines that are shown are the fitted values for participants 3SD (lightest blue), 2 SD (light blue) and 1 SD (blue) above the mean of the cheat count and participants 1 SD (dark blue), 2SD (darker blue) and 3 SD (black) below the mean of the cheat count.

#### Testing the predictive accuracy of the model

As the cognitive control regions were found to be most predictive of cheating, we used these predictors to test the prediction accuracy of our model. In order to do this, we used the trial level activation level in the ACC and left IFG, excluding the cheat count, obtained from the conjunction analysis and trained a multilevel logistic regression model, with random slopes and intercepts, on a training set (70% of the data). Subsequently, we tested the model on the left out 30% of the data. As the dependent variable, cheating, was imbalanced, we used two accuracy metrics that were insensitive to the class imbalance, namely the area under the curve (AUC) and the F1-score, which is the harmonic mean of the precision and recall. Statistical significance was estimated using permutation tests where the dependent variable (cheating) was permuted 5000 times and the classification metrics were estimated based on random permutations. We found that we were able to significantly predict cheating based on unseen data from the cognitive control network (AUC=76%, F1=89%, p<0.001).

### Individual differences in functional connectivity during decision-making

#### Connectivity within self-referential thinking network

The beta-series correlation analysis revealed that functional connectivity within nodes of the self-referential thinking network were more strongly connected for honest participants than for cheaters when making honest decisions. Specifically, correlations between honesty and functional connectivity were found between the PCC and left TPJ (r=0.51, p_adj_<0.05) and between PCC and MPFC (*r*=0.55, p_adj_<0.05; see Figure 7). No significant correlations between honesty and functional connectivity were found for cheated decisions. In addition, the correlation between honesty and functional connections between PCC and left TPJ and between PCC and MPFC during honest decisions were significantly different from the correlation during cheated decisions (both comparisons z > 2, *p_adj_*<0.005). Thus, the nodes within the self-referential thinking network, particularly between MPFC, left TPJ and PCC, seem to be more intimately connected to promote honesty particularly for honest participants, whereas when the connectivity between these nodes breaks down, honest participants tend to cheat.

**Figure 7.**
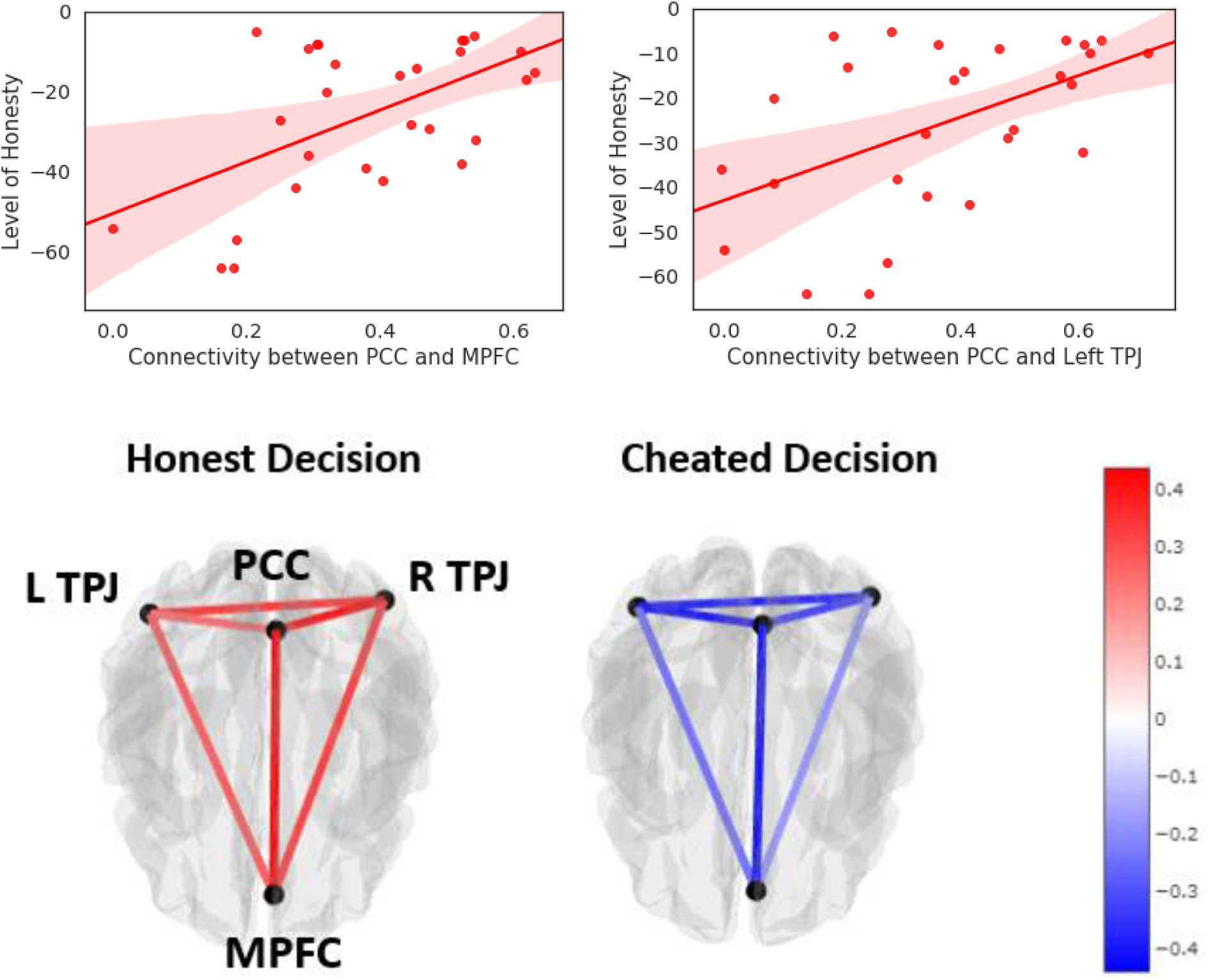
*Top row:* Correlation between level of honesty (reverse cheat count) and functional connectivity between PCC and MPFC and PCC and left TPJ. *Bottom row:* Connectome showing the correlation between level of honesty and the functional connectivity between within self-referential thinking network during cheated decisions (left) and honest decisions (right).

#### Classification of cheaters versus honest participants based on functional connectivity patterns

To test whether there is sufficient information in the connectivity patterns within the self-referential thinking network reported above to predict individual differences in honesty, a support vector classifier (Cox & Savoy, 2003; Mitchell, 2004) with linear kernel (C=1) was trained on the functional connectivity patterns of each participant to determine whether a participant was a cheater or an honest participant (categorized by median split). In order to avoid overfitting and inflated prediction accuracy (Vul, Harris, Winkielman, & Pashler, 2009) this was done using 8-fold cross validation (see Figure 8). Significance was estimated using permutation testing (N=5000). The classification analysis revealed that we could significantly classify an unseen participant as a cheater or an honest individual (Accuracy=75%, F1=71%, p<0.05).

**Figure 8.**
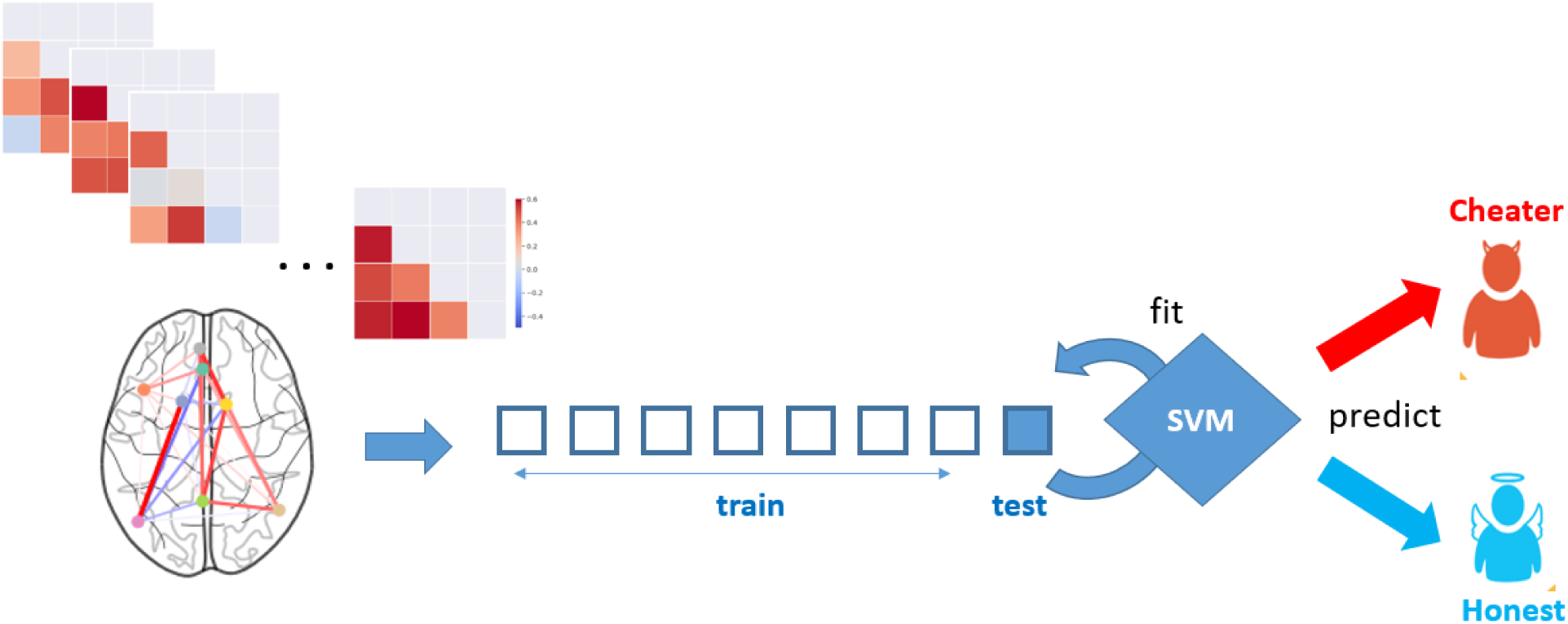
Using participants’ connectivity patterns within the self-referential thinking network during decision making to classify participants as cheaters or honest participants using support vector classifiers implemented with 8-fold cross-validation.

## Discussion

In this study we explored how neural mechanisms associated with reward anticipation, self-referential thinking and cognitive control determine the outcome of (dis)honest decisions. Using the newly developed Spot-The-Difference task to study trial-by-trial cheating behavior we found that the effect of cognitive control depends on a participant’s inclination to be honest or dishonest, in other words, on their moral default.

We found that more honest participants engaged in more self-referential thinking when exposed to the opportunity to cheat. Particularly, participants who were generally honest, exhibited higher activity in the self-referential thinking network comprised of the PCC, the bilateral TPJs and the MPFC. We confirmed that our results indeed reflect self-referential thinking processes by means of a conjunction analysis with Neurosynth data. Exploring the functional connectivity within the self-referential thinking network, we also found that more honest participants exhibited stronger positive connectivity during honest decisions between all nodes in this network, whereas this connectivity within the self-referential network broke down during cheated decisions. Collectively, these findings highlight the importance of our self-concept and related self-referential thinking processes in promoting honesty.

In line with previous research (Abe & Greene, 2014; Seuntjens et al., 2019), we found that cheaters exhibited stronger sensitivity to reward during decision-making. Our results revealed that all participants were anticipating reward and were sensitive to differences in magnitude of reward during the initial phase of the trial, where the potential reward for finding the differences between the two images is presented without moral conflict. However, cheaters, as compared to more honest participants, were more strongly driven by reward when making the decision whether to cheat or not. Specifically, cheaters exhibited higher neural activation in the Nacc, which is an area that has been consistently linked to reward anticipation (Ballard & Knutson, 2009; Knutson, Adams, Fong, & Hommer, 2001; Oldham et al., 2018), during the decision phase. Thus, whereas all participants are sensitive to differences in the magnitude of reward in the absence of moral conflict, particularly the cheaters are driven by the anticipation of reward when making the decisions to cheat.

Importantly, our study is the first to suggest that the function of cognitive control depends on a person’s moral default. Particularly, we found that for honest participants, more cognitive control, as represented by higher activity in the left IFG, was needed to cheat, whereas for participants who cheated frequently, response inhibition was needed in order to be honest. While honest participants needed cognitive control to overcome their inclination of being honest in order to cheat, cheaters had to exert control to override their greedy tendencies to be honest. Thus, our analyses indicated that the role of cognitive control depends on a person’s moral default.

In the literature there has been a debate between proponents of the “Will” hypothesis and the “Grace” hypothesis. Research supporting the “Will” hypothesis (Gino et al., 2011; Mead et al., 2009; Welsh & Ordonez, 2014) suggests cognitive control is needed to be honest. In direct opposition to this, another stream of research has accumulated evidence in favor of the “Grace” hypothesis (for meta-analyses, see Suchotzki, Verschuere, Van Bockstaele, Ben-Shakhar, & Crombez, 2017; Verschuere, Köbis, Bereby-Meyer, Rand, & Shalvi, 2018; Carparo, 2017; Spence et al., 2001; Greene & Paxton, 2009) advocating that cognitive control is required for dishonesty.

Our findings help reconcile this conflict as they suggest that people are distributed along a continuum from individuals who are generally honest to participants who can be considered cheaters. Participants on one side of the spectrum have a default inclination to be honest which is associated with more self-referential thinking when given the opportunity to cheat. In contrast, individuals on the other side of the spectrum have a default inclination for dishonesty and their decisions seem to be driven more strongly by rewards. In order to achieve and maintain a subjectively justifiable balance where one can occasionally profit from cheating but still maintain a positive self-image, people on both sides of the spectrum sometimes need to overcome their initial impulse and default behavior. A generally honest person will need to overcome the default of being honest in order to profit from cheating from time to time, whereas a cheater needs to inhibit the predominant selfish response in order to be honest and maintain their self-concept. Thus, it appears that the effect of cognitive control depends on our moral default. For honest people the “Grace” hypothesis applies: honesty results from the absence of temptation and response inhibition is needed to cheat. In contrast, for cheaters the predictions of the “Will” hypothesis apply and active resistance of temptation in form of inhibition is needed to be honest. Extending findings from cognitive psychology (MacLeod, 1991; Eriksen & Eriksen, 1974; Simon & Wolf, 1963) to the social/moral domain, our results suggest that cognitive control seems to serve the purpose to override our default behavior. Our study thus contributes to the reconciliation of the controversy on the role of cognitive control in moral decision making.

In addition, our findings also point to the importance of self-referential thinking processes and the maintenance of a positive self-concept. Whereas previous neuroimaging research has mainly focused on the role of cognitive control and reward sensitivity in cheating behavior, our study is the first to find neural evidence in favor of the self-concept maintenance theory (Mazar et al., 2008). Our results indicate that besides reward and control processes, self-referential thinking as represented by activation in the PCC, MPFC and bilateral TPJs, was engaged, particularly in honest participants, when they were tempted to cheat. In addition, regions in the self-referential thinking network were more strongly activated and functionally more strongly connected with each other when making honest decisions. Thus, our neural evidence suggests that when exposed to an opportunity to cheat particularly honest people do value their moral self-concept and its maintenance enough to forgo potential financial reward.

To examine the generalizability of our findings, we also tested the predictive power of the cognitive control regions on a trial-by-trial basis using cross-validation. We found that we could significantly predict with high accuracy on unseen data whether on a given trial participants would be honest or would cheat. Moreover, to assess whether connectivity patterns between the different networks contained relevant information about individual differences in honesty, we used support vector classifiers trained on participants’ connectivity patterns to discriminate cheaters from honest participants and found that we could indeed accurately classify whether a participant is a cheater or not. From the perspective of scientific rigor, cross-validation is a more conservative way to infer the presence of a brain-behavior relationship as compared to correlation or regression, as it is designed to protect against overfitting by testing the strength of the association in a new sample. This increases the probability of successful replication in future studies. Demonstrating this generalizability of our models may be an important first step in the development of useful neuroimaging-based biomarkers of dishonesty with real-world applicability.

In order to rule out alternative hypotheses we conducted several control analyses. First, in order to test whether neural differences during the decision phase were not driven by differences in levels of engagement with the task, we explored the neural processes during the visual search phase of each trial. As expected for a visual search task, we found that participants showed increased activation in areas related to visual and cognitive processing, working memory and navigation (see Appendix 7). More importantly, no significant differences in neural mechanisms during visual search were found between honest participants and cheaters. This eliminates the possibility that our neural findings were confounded by processes related to differences in engagement or effort during visual search. Second, we also conducted an exploratory factor analysis, which revealed that regions of interest used in our trial-by-trial and functional connectivity analyses indeed belonged to three separate networks that could be clearly identified as the control, reward and self-referential thinking network, respectively (see Appendix 8).

In reference to previous neuroimaging research on moral decision-making, our findings align with the early work using hypothetical moral dilemmas (Greene et al., 2004), instructed lying paradigms (Spence et al., 2001, Langleben et al., 2002) and also more recent work using the die-roll task (Greene & Paxton, 2009) in highlighting the importance of the cognitive control network, including areas such as the ACC and IFG, in moral decision-making. As stated above, our findings are also in line with those of Abe and Greene (2014) converging on the conclusion that a more sensitive and responsive reward network is associated with higher levels of cheating. With regard to neural processes linked to self-referential thinking, an fMRI study by Greene and colleagues (2001) found that a network of regions including the MPFC, PCC and bilateral TPJ were involved in making judgements about more personal as opposed to abstract hypothetical moral dilemmas, which they attributed to general emotional processes. More recently, a meta-analysis on neuroimaging research on moral decision-making, conducted by Lisofsky and colleagues (2014) reported that experimental deception paradigms that involved an identifiable victim and consequently perspective taking, were associated with increased activation in the right temporal parietal junction and the bilateral temporal pole, which have been associated consistently with Theory of Mind (ToM) processes (Bahnemann, Dziobek, Prehn, Wolf, & Heekeren, 2009) as compared to less interactive deception and cheating studies. Based on these findings, Lisofsky and colleagues argue that particularly in studies involving social interaction and an identifiable victim not only control processes but also perspective taking and moral reasoning processes are important. Our findings add to their conclusion by demonstrating that also in contexts without an identifiable victim, a similar network of regions, involving the TPJ but also the MPFC and PCC, is crucial in determining the outcome of moral decisions. This suggest that similar neural mechanisms may underlie self-referential thinking and perspective taking processes in the context of moral decision-making.

In contrast to our findings, a study by Garrett and colleagues (2016) showed that dishonesty escalates over time and is associated with diminished responsiveness in the amygdala, which they interpreted as reflecting reduced aversive emotional response to dishonesty. We did not find that dishonest behavior or the underlying brain networks changed over time. Garrett and colleagues used a paradigm with an identifiable victim which may explain why dishonesty increased with repetition in their study but was not observed in the Spot-the Difference task.

To conclude, we used a task that is the first to allow measuring cheating on the trial level in an fMRI environment. Using this novel task, we found that not only reward sensitivity but also the extent to which someone engages self-referential thinking processes determines whether someone is a cheater or tends to be honest most of the time. Importantly, we also found that the role of cognitive control on (dis)honesty depends on a person’s moral default. These findings may prove to be useful for developing interventions targeted at reducing cheating and dishonesty. Considering the huge economic losses caused by dishonest behavior, such as tax deception, music piracy or business scandals such as the Volkswagen emission fabrications, reducing dishonest behavior effectively is of great relevance to our economy and policy makers.

Taken together, we showed that the neural mechanisms engaged in (dis)honest decisions, ranging from neural activation in reward, self-referential thinking and control networks to functional connectivity patterns, differ fundamentally between honest and dishonest participants. Specifically, we found that cognitive control overrides a person’s moral default. Cognitive control allows honest people to cheat at times, whereas it enables cheaters to be honest. These insights contribute to a deeper understanding of neural correlates of individual differences in moral decision making. Future research may explore whether neural markers associated with dishonesty are also observable in more stable neural measures such as resting state functional connectivity or structural brain differences.

**Table 1.**
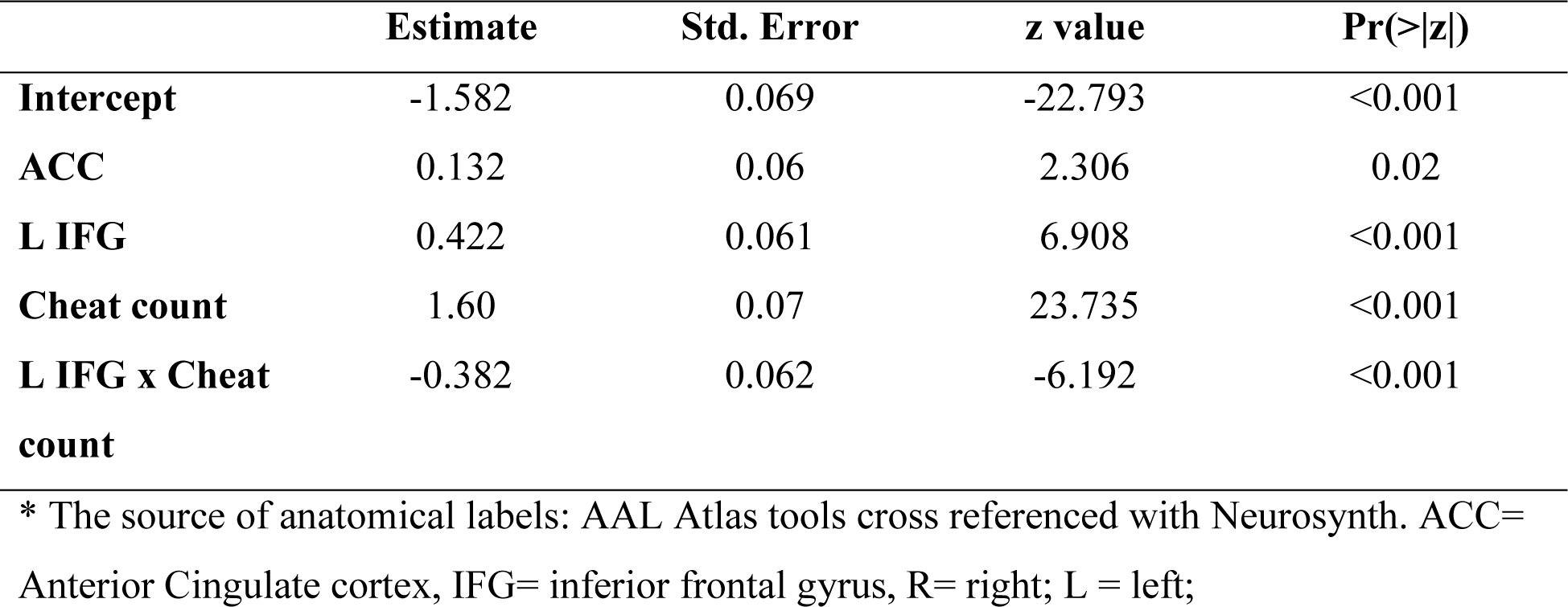
Multilevel Logistic Regression Model using the Cognitive Control Network to predict Cheating

## Appendices

## Appendix 1 Visual search task

To further increase the credibility of our cover story on brain processes underlying visual search, we also included the visual search task introduced by Treisman and Gelade (1980) at the beginning of our experiment. Specifically, participants were told that the experiment would start with a simple visual task and then proceed to visual searches in more complex visual stimuli in the second task. In this first task, the goal was to determine whether a specific target was present or absent. In each trial participants were presented with colored letters presented in random locations on the screen. If the target was present, then participants had to press the left mouse button as quickly as possible. If no target was present, then they had to press the right mouse button as quickly as possible. For this task, participants had to search for a green T. Participants were instructed to answer as quickly as possible while still being as accurate as possible. The task took approximately 5 minutes and was also completed in the scanner while localizer scans were obtained to ensure that scanning noise was audible, so participants would believe this task was indeed part of the study. This task will not be analysed as it was included solely for the purpose of increasing the credibility of our cover story.

**Figure S1.**
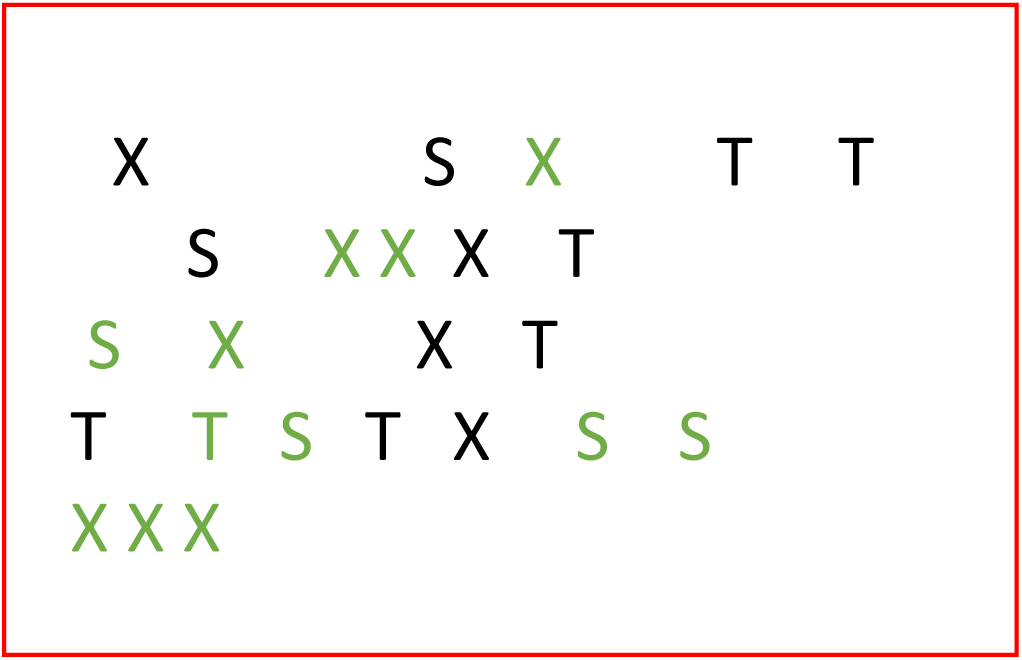
One trial of the simple visual search task. Participants have to indicate whether a green T is among the letters on the screen.

## Appendix 2 Validation of the picture set

Stimuli for the task consisted of 144 Spot-The-Difference image pairs that were downloaded from the Internet. Cartoon images of landscapes containing several objects were selected, to make them engaging and challenging enough for the participants. Landscapes were chosen as they generally satisfied the necessary criteria of containing several different objects, which made the task of spotting differences more challenging and engaging. The stimuli consist of pairs of images that are identical apart from a certain number (1-3) of differences that were created by the experimenter using Adobe Photoshop. Differences consisted of objects added to or removed from the landscape picture or changed colors of objects.

To make sure that participants would be able to find the differences between the images in a reasonable amount of time, we ran a pilot study on Amazon’s Mechanical Turk with 205 subjects using 180 pictures to test the difficulty to spot the differences between the images and to determine the optimal duration of picture presentation. Participants were presented with cartoon image pairs, presented horizontally next to each other, containing three differences and were asked to click on the differences identified in the image on the right hand side. They were given 15 seconds to make their response. Using the heatmap function provided by Qualtrics, regions of interest were defined around the locations of the differences in the image on the right hand side and response times for each of the clicks were recorded. This allowed us to test whether participants were able to find all differences in an image pair, which differences were particularly difficult to find, and how long it took to identify all differences. Based on the responses of these 205 participants, 36 image pairs that took too long or had differences that were too difficult or too easy, were removed, resulting in 144 images that took 92% participants less than 6s to find all three differences (M=5.4s, SD =1.5s).

## Appendix 3 Regions extracted for ROI analyses

Depicted here are the tables showing the regions extracted from Neurosynth.

**Table S1.** ToM and Cognitive Control masks link for download

| Network | Studies | Date of | Link to download |
| --- | --- | --- | --- |
| Self<br>Referential | 166 | 03.06.2019 | http://neurosynth.org/analyses/terms/self%20referential/ |
| Cognitive<br>Control | 598 | 03.06.2019 | http://neurosynth.org/analyses/terms/cognitive%20control/ |

## Appendix 4 Cluster statistics for the second-level results for cheatable vs non-cheatable trials

**Table S2.** Regions more activated during cheatable trials as compared to non-cheatable trials for honest participants as compared to cheaters

| Region | cluster_id | peak_x | peak_y | peak_z | peak_value | volume_mm |
| --- | --- | --- | --- | --- | --- | --- |
| PCC | 1 | -9 | -57 | 23.79 | 492.667 | 10014 |
| R TPJ | 2 | 45 | -60 | 23.79 | 445.751 | 4138 |
| Hippocampus | 3 | 24 | -18 | -18.33 | 440.276 | 3632 |
| (v)MPFC | 4 | -6 | 54 | -4.29 | 388.299 | 3538 |
| Cerebellum | 5 | 0 | -54 | -60.45 | 379.483 | 3222 |
| MFG | 6 | -30 | 24 | 44.85 | 407.816 | 3032 |
| Cerebellum | 7 | -27 | -48 | -25.35 | 429.123 | 2811 |
| Left Frontal Pole | 8 | -18 | 39 | 44.85 | 421.571 | 2337 |
| MPFC | 9 | -6 | 30 | 6.24 | 39.304 | 2053 |
| L TPJ | 10 | -45 | -69 | 23.79 | 38.957 | 1674 |
| R Postcentral |  |  |  |  |  |  |
| Gyrus | 11 | 30 | -42 | 65.91 | 382.644 | 1547 |
| R Supramarginal |  |  |  |  |  |  |
| Gyrus | 12 | 66 | -30 | 27.3 | 448.577 | 1263 |
| L Supp motor area | 13 | 0 | 0 | 48.36 | 365.033 | 1232 |
| L C | 14 | -18 | -42 | -49.92 | 411.611 | 1105 |
| R Cerebelum 4 5 | 15 | 12 | -45 | -11.31 | 429.168 | 1105 |
| R Hippocampus | 16 | 21 | -39 | 6.24 | 466.645 | 1105 |
| L OFC | 17 | -42 | 36 | -14.82 | 431.246 | 1042 |

## Appendix 5 Cluster statistics for the second-level results for cheated vs honest decisions

**Table S3.** Regions more activated during honest decisions as compared to cheated decisions for cheaters than for honest participants

| Region | cluster_id | peak_x | peak_y | peak_z | peak_value | volume_mm |
| --- | --- | --- | --- | --- | --- | --- |
| L IFG | 1 | -46 | 21 | -5 | 41 | 4156 |
| R ACC | 2 | 7 | 36 | 21 | 384 | 2797 |
| L ACC | 3 | -7 | 41 | 7 | 450 | 1922 |
| R Insula | 4 | 37 | 26 | -6 | 387 | 762 |
| L Frontal Pole | 5 | -34 | 62 | 5 | 476 | 704 |
| L Supp Motor Area | 6 | -11 | 23 | 63 | 398 | 639 |
| L Nacc | 7 | -14 | 19 | -7 | 372 | 326 |
| L SFG | 8 | -4 | 20 | 43 | 356 | 272 |
| R Cingulate Gyrus | 9 | 1 | -28 | 29 | 330 | 237 |
| R Angular Gyrus | 10 | 54 | -51 | 45 | 331 | 200 |

## Appendix 6 Cluster statistics for the second-level results of the parametric modulation analysis for the level of reward

**Table S4.** Regions parametrically modulated by level of reward during the level of difficulty phase of the Spot-The-Difference task

| Region | cluster_id | peak_x | peak_y | peak_z | peak_value | volume_mm |
| --- | --- | --- | --- | --- | --- | --- |
| Left Cuneus Cortex | 1 | -9 | -78 | 16.77 | 546 | 1611 |
| R Nacc | 2 | 12 | 12 | -0.78 | 493 | 1232 |
| L Nacc | 3 | -21 | 15 | -0.78 | 47 | 568 |
| L Cuneus | 4 | -6 | -96 | 27.3 | 414 | 315 |

## Appendix 7 Levels of engagement during visual search

In order to test whether our findings may be confounded by different levels of engagement during the visual search phase, we tested whether there were differences in neural activation during the visual search phase between more honest participants and cheaters. First, we ran a univariate analysis in which we contrasted neural activity during the visual search against baseline activation. The analysis revealed that a big cluster in the visual cortex showed higher activation during search as compared to baseline activation, which is expected as participants were engaged in visual search. In addition, several regions related to working memory, cognitive processing and navigation, such as the dmPFC and the MFG were more strongly activated during visual search (see Appendix for table with cluster statistics). To explore whether there are individual differences in level of engagement during visual search, participants’ cheat count was added as a group level covariate. The whole brain analysis revealed that there are no significant differences between more honest participants and cheaters during the visual search phase. In addition, we also tested whether differences in neural activation during visual search between cheatable and non-cheatable trials were more strongly expressed in cheaters or honest participants. In order to do so, a univariate analysis was run in which we contrasted neural activation during visual search in cheatable trials against activation during visual search in non-cheatable trials. Again, these contrast maps were then correlated with cheat count on the group level. The whole brain analysis did not reveal any significant effects. These findings suggest that there are no significant differences in level of engagement or motivation during visual search between more honest participants and cheaters.

**Figure S2.**
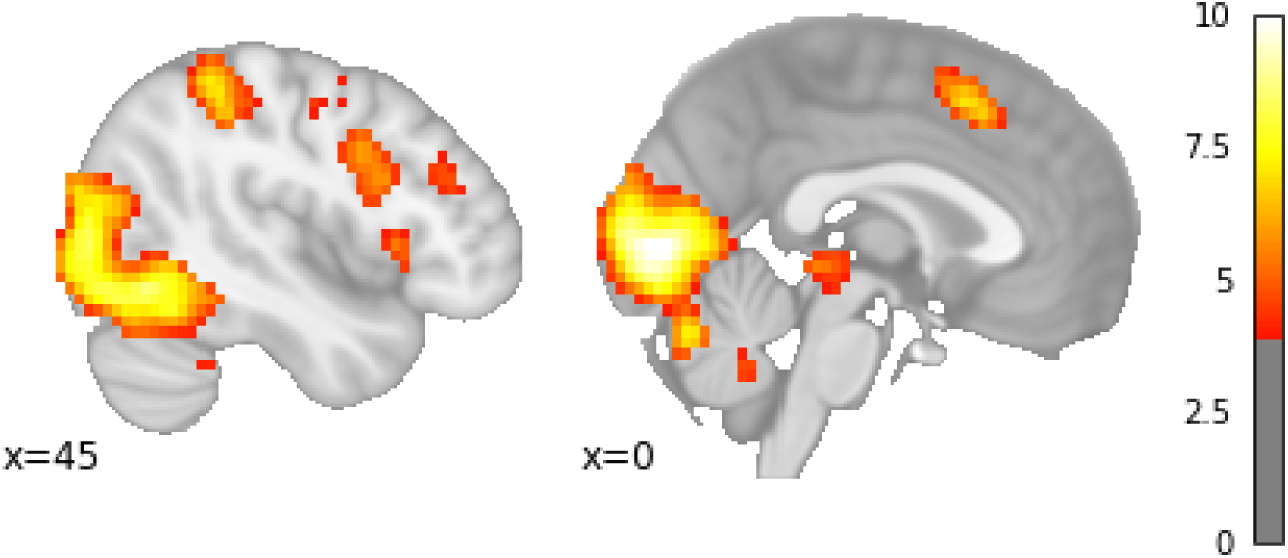
The visual cortex, dMPFC and left and right dlPFC are more activated during visual search as compared to baseline

**Table S5.**
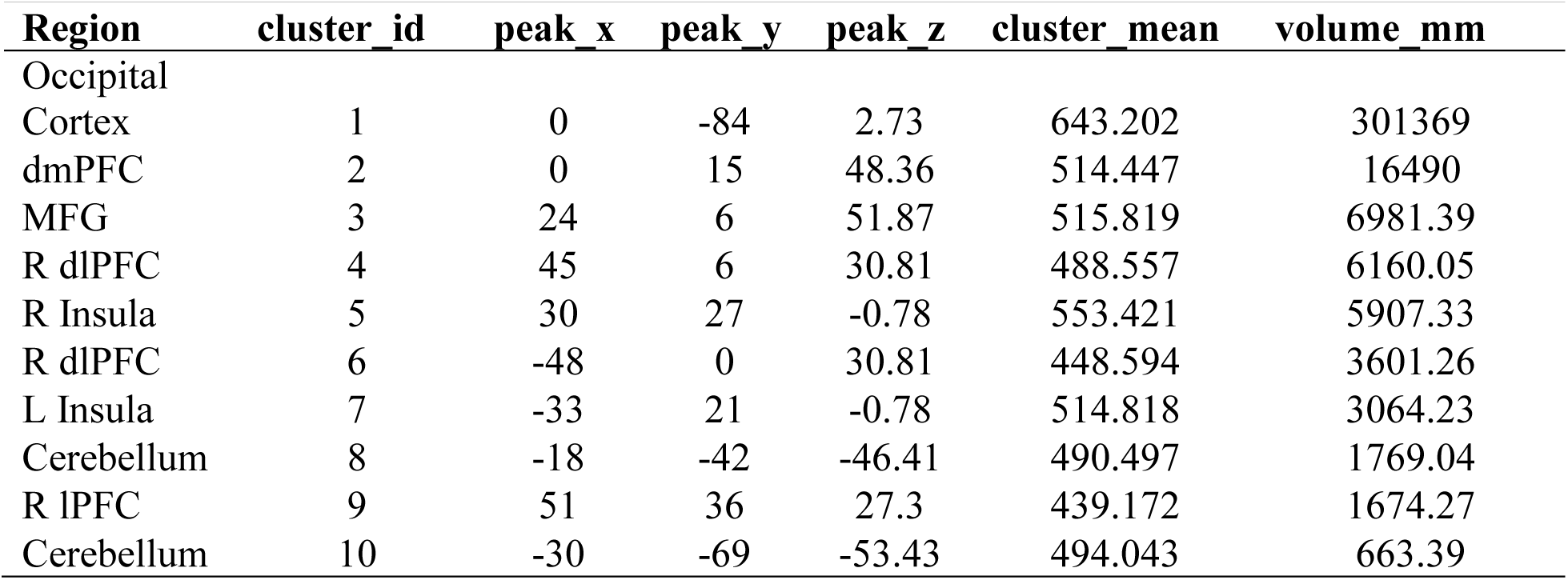
Regions more activated during visual search as compared to during rest

## Appendix 8 Factor analysis to confirm validity of networks

To test whether the regions we are analyzing indeed belong to three separate networks, we conducted an exploratory factor analysis with promax rotation (Hendrickson & White, 1964), which is an oblique rotation method which allows for correlation between latent factors. Specifically, the goal of this factor analysis was to determine the most important latent factors underlying all the regions resulting from our conjunction analyses, namely the left IFG and ACC (cognitive control network), the PCC, bilateral TPJs and MPFC (self-referential network), and the bilateral Nacc (reward network). We used the single trial activations obtained as explained above by fitting a model that includes a separate regressor for each trial from each of the regions as input for the factor analysis. Before conducting the factor analysis, we first checked whether the regions intercorrelated at all using Bartlett’s test of sphericity which tests the observed correlation matrix against the identity matrix. Bartlett’s indicated that the null hypothesis can be rejected and there is significant correlation between variables justifying a factor analysis (χ2 =10582, p<0.001). In addition, the Kaiser-Meyer-Olkin (KMO) test was conducted which determines the adequacy of the observed variables by estimating the proportion of variance among all the observed variables. The KMO test revealed an overall estimate of 0.69 which indicates that the observed variables are adequate for a factor analysis. Next, we determined the number of factors with the help of the Kaiser criterion (choosing factors with an eigenvalue >1). This resulted in three latent factors, where the first factor represented the self-referential thinking network with the bilateral TPJs, PCC and the MPFC loading highly on this factor. The second factor clearly represents the reward network as only the bilateral Nacc show high factor loadings. Lastly, the third factor clearly represents the cognitive control network as only the ACC and the left IFG load highly on this component. This exploratory factor analysis clearly indicates that the regions of interest used in our trial-by-trial and functional connectivity analysis indeed belong to three separate networks.

**Figure S3.**
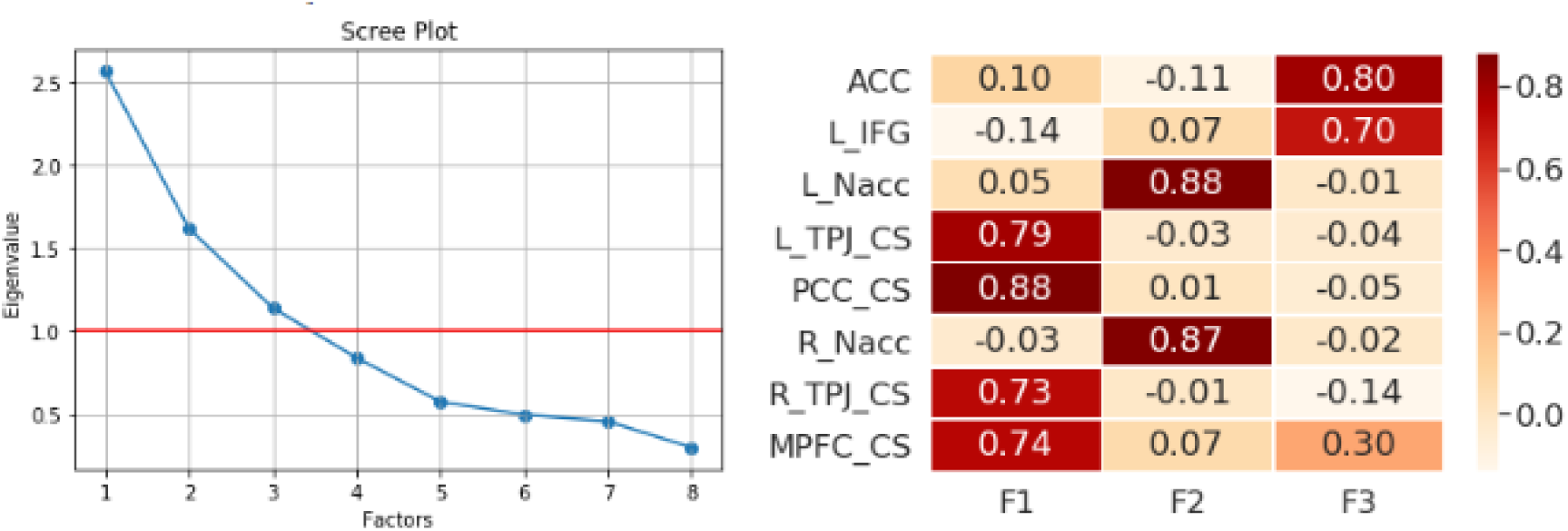
Left: Scree plot showing the the eigenvalues for each factor. Right: the loadings for each of the factors

## Appendix 9 Classifying cheaters versus honest participants

Due to the fact that we found that we could classifiy cheaters and honest participants based on the functional connectivity patterns during decision-making, we wanted to see whether average activation within a subject in the ROIs from the three networks of interest (cogntive control, reward & self referential thinking) could be used to classify participants as cheaters or honest participants (categorized by median split). In order to do this we average the trial by trial estiamtes within participants resulting in one observation for each subject, which represents the average activation in each ROI across the whole task. In order to test this we employed a support vector classifier (Cox & Savoy, 2003; Mitchell, 2004) with linear kernel (C=1) was trained on average activations in the ROIs of each participant to determine whether a participant was a cheater or an honest participant (categorized by median split). In order to avoid overfitting and inflated prediction accuracy (Vul et al., 2009) this was done using 8-fold cross validation. Significance was estimated using permutation testing (N=5000). The classification analysis revealed that we could significantly classify an unseen participant as a cheater or an honest participant based on the average activation in the ROIs (F1=70%, AUC=77%, p<0.05). Using activations from honest trials only an even higher classification accuracy was found (AUC=0.84, p<0.05). Classification was not significant using cheated trials only.

